# Mechanical loading induces distinct and shared responses in endothelial and muscle cells and reveals exercise-like molecular profiles

**DOI:** 10.1101/2025.06.27.661962

**Authors:** Sakari Mäntyselkä, Erik Niemi, Laura Ylä-Outinen, Kalle Kolari, Liina Uusitalo-Kylmälä, Alfredo Ortega-Alonso, Roosa-Maria Liimatainen, Vasco Fachada, Perttu Permi, Elina Kalenius, Juha J Hulmi, Riikka Kivelä

## Abstract

Skeletal muscles and blood vessels are continuously exposed to mechanical forces, particularly during exercise. We subjected human endothelial and skeletal muscle cells to cyclic mechanical stretch to mimic exercise and investigated acute molecular responses. Mechanical loading elicited both shared and cell type-specific alterations in transcriptomic and metabolomic profiles, several of which mirrored changes observed in vivo following exercise. Both cell types released acetate in response to mechanical loading, at least partly via reactive oxygen species -dependent mechanism. Interestingly, transcriptomic changes occurred in opposite directions in endothelial and muscle cells. For example, genes associated with the electron transport chain were repressed in endothelial cells but upregulated in skeletal muscle cells. In endothelial cells mechanical loading remodelled intercellular junctions, promoted a transcriptomic shift indicative of increased barrier integrity and attenuated proliferation. Metabolic changes were more pronounced in endothelial cells, which exhibited increased serine biosynthesis from glucose, as demonstrated by ^13^C-(U)-glucose tracing. Targeting phosphoglycerate dehydrogenase (PHGDH), a key enzyme in the serine synthesis pathway, underscored the role of serine biosynthesis in endothelial cell anabolism. These findings suggest that mechanical loading recapitulates several exercise-induced effects in endothelial and muscle cells, and highlights a potential link between mechanical stimuli, serine synthesis, and endothelial cell quiescence.

## 1 Introduction

Skeletal muscle is a highly vascularized tissue and myofibers and endothelial cells together make up ∼70% of the total nuclei count in skeletal muscle tissue (1, 2). Skeletal muscle is a highly mechano-sensitive tissue, responding to increases or decreases in mechanical load by changes in function and morphology, and it is well established that increased mechanical loading, by e.g. resistance exercise, is a crucial factor in exercise-induced skeletal muscle hypertrophy (3). Furthermore, during exercise, up to 80-90% of cardiac output is directed to contracting skeletal muscle (4) which also elicits increased mechanical loads such as shear stress and circumferential wall stress (stretch) on endothelial cells lining the blood vessels, and these hemodynamic forces play a central role in the exercise-induced changes in endothelial cell phenotype (5). However, it is difficult to distinguish the effects of mechanical stretch alone in *in vivo* settings, which is why *in vitro* contraction and stretching models have been used to mimic these stimuli (6, 7). Previously, it has been demonstrated that cyclic mechanical loading (i.e. cyclic stretching) induces a wide variety of molecular and morphological responses in endothelial [reviewed in (8)] and muscle cells [reviewed in (9)]. For example, mechanical loading has been found to activate the mechanistic target of rapamycin complex 1 (mTORC1) (10) and to induce highly similar gene expression changes in murine myofiber-like multinucleated cells (myotubes) as resistance exercise *in vivo* in myofibers (11). Still, studies using human muscle cells are needed (7). Endothelial cells, in turn, have been shown to respond to mechanical stretch in various ways, such as increased migration and changes in orientation, secretion of angiocrines, increased proliferation, and activation of angiogenesis-related pathways (8, 12, 13), which also occur in response to exercise and increased blood flow *in vivo* (14, 15). On the other hand, mechanical stretch of endothelial cells has also been reported to increase apoptosis, inflammation, and formation of reactive oxygen species (ROS), depending on whether the magnitude of the stretch is categorized as physiological (∼10%) or pathological (over 20%) (8). However, there are only a few studies (12, 16, 17) investigating the remodeling of the global molecular landscape in response to mechanical loading, and none in comparing the responses in different cell types.

Skeletal muscle responses to mechanical loading *in vivo* have mostly been examined using mixed muscle samples, which lack cell-specificity as skeletal muscle contains over 10 different cell types (1), and a recent study investigating the acute transcriptional responses to exercise in mice demonstrated that endothelial cells show a distinct gene expression response from muscle cells (18). *In vitro* models allow us to study the transcriptomic and metabolic responses in a cell type-specific manner. Previously, similar mechanical stretching of liver endothelial cells during vasodilation was demonstrated to induce angiocrine signals that support liver growth and survival, and potentially contribute to liver regeneration (13). It is also recently acknowledged that endothelial and muscle cell cross-talk is essential in regulating the development, maintenance, and repair of muscle tissue [reviewed in (2)]. However, despite these recent findings, according to our knowledge, there are no studies in which the responses elicited by mechanical loading alone in muscle and endothelial cells have been compared. Exercise induces many metabolic changes, such as increased glycolysis and oxidative phosphorylation at the whole-muscle level (19). However, little is known about how exercise or mechanical loading *per se* affects other metabolic pathways, such as those branching from glycolysis (e.g., the serine synthesis pathway), and their role in exercise-induced responses.

Here we explored how mechanical loading affects the transcriptional landscape and metabolomics of two different human endothelial cells, and primary human skeletal muscle cells, to mimic exercise on-a-dish and to study cell type-specific responses. Molecular responses resembled changes typically observed after exercise *in vivo*, however, many pathways were changed in opposite directions in endothelial cells and muscle cells, demonstrating highly cell type-specific responses.

## 2 Materials and methods

### 2.1 Endothelial and muscle cells

We used human umbilical vein endothelial cells (HUVEC, Cat# C-12200, PromoCell), human cardiac microvascular endothelial cells (HCMEC, Cat# C-12285, PromoCell), and primary human skeletal muscle (HSkM) cells. The latter were obtained as previously (20) from two healthy 25-34-year-old volunteers (1 male and 1 female). Vastus lateralis muscle biopsies were obtained under local anesthesia with the Bergström biopsy needle. The human study (21) was conducted in accordance with the Declaration of Helsinki and was approved as part of a larger TraDeRe-study by the Ethics Committee of the University of Jyväskylä (857/13.00.04.00/2021). All participants provided written informed consent for inclusion prior to participating in the study.

### 2.2 Cell culture

The HUVEC and HCMEC cells were cultured in endothelial cell growth medium (EGM) (Cat# C-22010, PromoCell) supplemented with EGM SupplementMix (Cat# C-39215, PromoCell) and 1% (vol/vol) of 1000 U/ml penicillin-streptomycin (P/S, Cat# 11548876, Gibco). HSkM cells were cultured in growth medium (GM) consisting of high glucose (4.5 g/l) Dulbecco’s Modified Eagle Medium (DMEM, Cat# 10313021, Gibco) supplemented with 20% (vol/vol) fetal bovine serum (FBS, Cat# 10270106, Gibco), 1% (vol/vol) of 1000 U/ml P/S and 1% (vol/vol) 200 mM L-glutamine (Cat# 11500626, Gibco).

After reaching over 80% confluency in culture flasks, the cells were seeded (∼20,000 cells/cm^2^) on polydimethylsiloxane (PDMS) chambers (Cat# STB-CH-04, STREX). PDMS chambers were pre-coated with FBS for 24 h and with 0.2% gelatin for an additional 24 h. The PDMS chambers were placed in 150 mm diameter Petri dishes, two chambers per dish. Cells were grown on PDMS chambers until reaching over 80% confluency. After that, HUVEC and HCMEC cells were mechanically loaded, as described in the next section. In the case of HSkM cells, at 80% confluency the GM was replaced with differentiation medium (DM) containing high glucose DMEM, 2% (vol/vol) horse serum (HS, Cat# 26050088, Thermo Fisher Scientific), 1% P/S, 1% L-glutamine and 1% (vol/vol) Insulin-Transferrin-Selenium-X supplement (ITS-X, Cat# 51500056, Gibco). To induce myoblast fusion into myotubes, the cells were cultured in DM for 4-5 days and DM was changed every 48 h. Cells were cultured at 37 °C in a 5% CO_2_ atmosphere all the time.

### 2.3 Mechanical loading set-up

Mechanical loading of cultured cells was achieved with a commercial cell culture pacing system (C-Pace EM 100, IonOptix) paired with a motor-powered mechanical stimulation unit (C-Stretch 100, IonOptix). The C-Stretch mechanical stimulation unit consists of a motor-driven tray that causes uniaxial elongation of the PDMS chambers inserted in the device and exposes cells, that are attached to the bottom of the chamber, to mechanical load.

As mentioned, cells were seeded onto 12 PDMS chambers for each stretch experiment, with 6 chambers loaded into the C-Stretch unit in series, and 6 chambers left as static controls. The stretch unit was thoroughly cleaned with 70% EtOH before use. Fresh media was changed for all chambers prior to starting the experiment. The stretch unit was then placed in an incubator with conditions of 37 °C and 5% CO_2_ atmosphere along with the 6 control chambers. A cyclic mechanical stretch with a frequency of 1 Hz and 12% elongation was applied for 5 h. For each one-second cycle, a trapezoidal waveform of 0.4 s stretch, 0.1 s hold, 0.4 s return, and 0.1 s hold, was used to keep the velocity of the deformation of the silicone chamber at a moderate speed.

### 2.4 Lactate dehydrogenase activity assay

Lactate dehydrogenase activity (LDH) was measured from conditioned media as an indicator of possible membrane damage and cell death (22). The analysis was performed as previously (20) using an LDH Activity Assay Kit (Cat# 981906, Thermo Fisher Scientific).

### 2.5 RNA extraction and analyses

#### 2.5.1 RNA isolation, library preparation and sequencing

RNA was extracted using a commercial column-based RNA isolation kit (NucleoSpin® RNA, Cat# 740955.250, Macherey-Nagel) immediately after the 5-hour mechanical loading. The PDMS chambers were washed with PBS, and cells were lysed by pipetting back and forth vigorously with lysis solution (350 µl of guanidium thiocyanate lysis buffer + 3.5 µl of β-mercaptoethanol). Extraction was then continued according to the kit’s instructions. An additional centrifugation step of 30 s at 11,000 x g was included at the end to ensure complete removal of the wash buffer. Bound RNA was then eluted into a volume of 60 µl of RNase-free water. The collected elution was then aliquoted and samples were immediately transferred to - 80 °C. RNA quality control, library preparation and sample sequencing were performed at Novogene (Cambridge, United Kingdom), utilizing an Illumina NovaSeq 6000 platform, producing 150bp paired-end reads. Filtering of raw sequence data was also conducted by Novogene in the following way: (1) removing reads containing adapters, (2) removing reads containing >10% of unsolvable bases and (3) removing reads containing >50% of low quality bases (Qphred ≤ 5). Filtered RNA sequencing data is available in the Gene Expression Omnibus (GEO) data repository (accession number: GSE288064).

#### 2.5.2 Read alignment, normalization and filtering

Reads were aligned to the human genome (build GRCh38.95) using the Bowtie2 software (23), with alignment rates >80% for all the samples. The aligned and mapped reads were later annotated and quantified using the Rsubread package in R/Bioconductor software (24), with the human GRCh38.110 genome build as annotation reference. The obtained gene counts were transformed to counts-per-million (CPM) using the edgeR package in the R/Bioconductor software (25). Genes with a CPM < 1 in three samples or more in each group (50%), were subsequently filtered out as described in (26). The resulting expression data were finally normalized using the trimmed mean of M-values (TMM) algorithm (27).

#### 2.5.3 Differential gene expression and pathway analyses

Differential gene expression (DGE) analyses were implemented through the R/Bioconductor limma-voom pipeline (28). The voom-function of the limma package was initially executed to format the data appropriately (29), after which empirical Bayesian statistics were computed to rank genes based on their evidence for differential expression. To account for type I errors induced by multiple testing, the Benjamini-Hochberg method was used to adjust the tests’ *p-*values, retrieving an adjusted *p*-value (adj. *p*). Genes with a log_2_FC ≥ 0.5 and adj. *p* ≤ 0.05 were regarded as differentially expressed. For subsequent Gene Set Enrichment Analysis (GSEA) genes were ranked in an unbiased way by multiplying the log_2_FC by the -log10 of the adj*. p* to consider both the magnitude and consistency of the change in gene expression. GSEA was performed with the clusterProfiler R-package (30) with Gene Ontology: Biological Process (GO:BP), Reactome, and the Kyoto Encyclopedia of Genes and Genomes (KEGG) serving as reference databases. Ontologies and pathways were considered significantly enriched if their corresponding tests yielded an adj. p-value <0.05. In addition, gene set overrepresentation analysis (ORA) of the significantly up- or downregulated genes (log_2_FC ≥ 0.5 & adj. *p* ≤ 0.05) was conducted using the ShinyGO web-application v0.81 (https://bioinformatics.sdstate.edu/go/) (31) from GO:BP pathways. Pathway minimum size was set to 2 and maximum to 5000 and the FDR cutoff for significant pathway enrichment was set to 0.05. All analyses implemented in R/Bioconductor included the core software version 4.3.1 with the R-studio interface version 2024.04.2.

### 2.6 Immunocytochemical imaging and EdU analysis

Effects of mechanical loading on HUVEC morphology was assessed by immunocytochemical staining of endothelial membrane proteins and imaging with confocal microscopy. Cells were mechanically loaded as described in section 2.3 and stained and imaged directly on the PDMS chambers. Cells were first washed twice with PBS and fixed with 4% PFA for 15min. Primary antibodies for VE-Cadherin (Cat# 2500S, Cell Signaling), Claudin 5 (Cat# 34-1600, Invitrogen) and ZO-1 (Cat# 40-2200, Invitrogen) were diluted in PBS with 1% normal goat serum (NGS) and 1% bovine serum albumin (BSA) in concentrations of 1:400, 1:200 and 1:200, respectively, and incubated overnight in +4°C. Cells were then washed three times with PBS and cells were incubated in secondary antibody (Cat# A11008, Invitrogen) diluted 1:300 in PBS for 1h in RT, protected from light. After 1h, cells were washed with PBS and incubated in DAPI solution in PBS (0.5 µg/ml) for 15min in RT to visualize nuclei. Cells were finally washed three times with PBS and imaged with confocal microscope (Zeiss LSM700).

Effects of mechanical loading on proliferation of HUVECs were assessed both by microscopy and flow cytometry. For both analyses, the cells were mechanically loaded as described earlier and incubated in 5-ethynyl-20-deoxyuridine (EdU) labeling solution (Cat# C10339, Invitrogen) during the mechanical loading (5h), plus an additional 2h after cessation of the loading period. For the microscopy analysis, cells were fixed with 4% PFA for 15 min, washed with 3% bovine serum albumin (BSA) in PBS, and permeabilized with 0.5% Triton X-100 in PBS for 20 min RT, and the Click reaction was done according to the manufacturer’s instructions. Before imaging, DAPI (0.5 µg/ml) was added. Cells were imaged directly from PDMS chambers. Quantification of EdU-positive nuclei was performed with an in-house built Fiji script (https://github.com/seiryoku-zenyo/simple_ratio_DAPIvsPositive-nuclei/tree/main). Shortly, image data was segmented using a deep learning algorithm, separately for DAPI-positive nuclei and EdU-positive nuclei. Next, the ratio between DAPI and EdU nuclei was counted. The data is presented as a fraction (%) of EdU-positive nuclei out of total (DAPI) nuclei.

For flowcytometry analysis, the Click-iT™ EdU Alexa Fluor™ 647 Flow Cytometry Assay Kit (Cat# C10424, Invitrogen) was used. HUVECs were detached after the EdU incubation, and two PDMS chambers were pooled as one sample to ensure enough cells for the flow cytometer. Fixing, washing, and staining of the cells were performed following the kit instructions. EdU analysis was performed with CytoFLEX S Flow Cytometer using red 638 nm laser with 712/25 BandPass filter and Cytexpert software 2.4.0.28. Quality control was performed using CytoFLEX Ready-To-Use Daily QC Fluorospheres, and an unstained HUVEC pool was used as a compensation control. 10000 datapoints were collected. Cell debris and determination of singlets with SSC and FSC were performed. Analysis of EdU^+^ and EdU^-^ signals was done utilizing histograms.

Click-iT EdU imaging kit (Cat# C10339, Invitrogen) was also used to determine the effect of PHGDH overexpression (see section 2.13) on HUVEC proliferation. Cells were cultured on 13 mm diameter coverslips (10 000 cell/cm^2^) coated with 0.2% gelatin for 24 h, after which cells were incubated with EdU labeling solution for 5 h. Cells were fixed, washed, and permeabilized as above. Click reaction buffer was prepared as described by the manufacturer, added to the cells, and incubated for 30 min protected from light. DAPI was used to detect all the nuclei. Cells were imaged with a confocal microscope (Zeiss LSM700), and here the number of stained nuclei was calculated manually with Fiji. The data is presented as a fraction (%) of EdU-positive nuclei out of total (DAPI) nuclei.

Effects of PHGDH inhibitors (see section 2.11) on cell division capacity were evaluated using a Click-iT EdU imaging kit (Cat# C10338, Invitrogen). HUVEC cells were plated onto a 96-well plate in Endothelial cell GM and let to grow in subconfluent culture for 24 h. Exposures were added to the wells, four parallel wells for each condition, and eight parallel wells for the control condition (DMSO vehicle). After 20 h incubation of exposures, EdU was added for 4 h. Cells were imaged using a Nikon A1R confocal microscope. DAPI was used to detect all the nuclei. The microscope was automated to find optimal focus in each well, image every well, and take 2-9 views per well so that at least 300 nuclei were imaged. Quantification of EdU-positive nuclei was performed with an in-house built Fiji script as above.

### 2.7 Carbon-14 labeling

For evaluation of relative protein synthesis and glucose-incorporation the following radioisotope tracers were used: ^14^C-[1]-L-phenylalanine (0.1 mCi/mL, Cat# NEC859S050UC, Revvity); ^14^C-[U]-L-valine (271.0 µCi/mmol, Cat# NEC291EU050UC, PerkinElmer); and ^14^C-[U]-glucose (0.2 mCi/mL, Cat# NEC042V250UC, PerkinElmer). Only one tracer per experiment was used, and the dilution of the tracers was 1 µL/mL (vol/vol) in the culture media. Two milliliters of culture media were used per well of a 6-well plate, and one milliliter per stretching chamber

When protein synthesis was measured during and after mechanical loading, tracer was added to the medium just before the initiation of the 5 h mechanical loading. After the loading, HUVECs were incubated for an additional 20 h in the medium, and then proteins were extracted (see section 2.14). HUVECs for PHGDH inhibitor experiments were seeded onto 0.2% gelatin-coated 6-well plates at a density of 12,500 cells/cm². The cell population was expanded for 2 days, and the inhibition experiments were initiated by changing the media to experimental media containing 25 µM of inhibitor and a radioisotope tracer. After incubating cells for 48 h with inhibitor and tracer, proteins, RNA, and lipids were extracted, and liquid scintillation counting was conducted as previously (20).

### 2.8 Meta-analysis of skeletal muscle response to exercise (MetaMEx)

We examined gene expression of the *Endothelial cell development* pathway (GO:0001885, 72 genes, list checked on 10.06.2025) in response to acute aerobic exercise using the MetaMEx browser-based tool (https://www.metamex.eu/app/metamex) (32). All the other settings were kept as a default, except that only the healthy population was included.

### 2.9 Metabolite extraction and analyses

#### 2.9.1 ^1^H NMR spectroscopy metabolomics

Metabolites from the media samples were extracted as previously (20, 33). Cell-free medium samples were also included as a reference to assess the consumption and release of metabolites, as previously (33, 34). Dried samples were reconstituted with 200 µL of sodium phosphate buffer (150 mM, pH = 7.4) in 99.8% deuterium oxide (Acros Organic; Thermo Fisher Scientific) containing 0.5 mM 3-(trimethylsilyl)propanesulfonic-d_6_ acid sodium salt (DSS-d_6_; IS-2 Internal Standard, Chenomx Inc.) and transferred to round-bottom 3-mm OD NMR tubes (Cat# 341-PP-7, Wilmad). Data collection was carried out at 25 °C using the Bruker AVANCE III HD NMR spectrometer, operating at 800 MHz ^1^H frequency equipped with a cryogenically cooled ^1^H, ^13^C, ^15^N triple-resonance probe head (Bruker Corporation). The ^1^H nuclear magnetic resonance (^1^H NMR) spectra were collected as previously described (33). ^1^H NMR spectra were processed, and metabolites were identified and quantified with the Chenomx 10.0 software (Chenomx).

#### 2.9.2 UHPLC-MS/MS targeted metabolomics

Intracellular metabolites were collected and extracted as previously (20) with minor modifications. In brief, 200 µL of ice-cold 80% methanol was added to each stretching chamber, and metabolites from three chambers were pooled for one sample. Dried samples were reconstituted with 40 µl of 50-50 acetonitrile-water mixture (vol/vol) before analysis. Metabolites were analyzed as previously (20). Briefly, metabolites were separated with an Atlantis Premier BEH Z-HILIC UHPLC column (1.7 µm, 2.1 x 100 mm) which was used with an Agilent 1290 Infinity ultra-high-performance liquid chromatography (UHPLC) system. *Mobile phase A* was 15 mM ammonium bicarbonate (pH = 9.0) in water, and *mobile phase B* was 15 mM ammonium bicarbonate (pH = 9.0) in 90% acetonitrile—10% water. The flow rate was 0.7 mL/min, the column oven temperature was 40 °C, and the injection volume was 3 µl. The following gradient elution was used: the initial concentration of 90% *B* was held for 1 min and then was linearly decreased to 65% *B* over the next 5 min and was held for 1 min at 65% *B*. Then *B*% was increased to 90% over the next 1 min and was held 2 min at 90% *B* before next injection. The UHPLC system was coupled with an Agilent 6460 triple quadrupole mass spectrometer operated with dynamic multiple reaction monitoring mode (dMRM) as previously described (20). A mixture of target metabolites (concentrations between 10 and 50 µM) was injected eight times at the beginning of the analysis and after every 6 to 8 samples for quality control and metabolite identification purposes.

Acetate was analyzed from media samples using the above-described chromatographic method and the UHPLC-MS instrument which was operated on negative polarization. For acetate, selected ion monitoring [e.g., pseudo-MRM (35)] of *m/z* 59 was done. The following parameters were used for the selected ion monitoring experiment: fragmentor: 80; collision energy: 1; cell accelerator voltage: 4. The other ion source parameters were the same as previously reported (20). Retention time and linearity of the signal were tested using 10, 20, 40, 80, and 160 µM acetate standards that were prepared using sodium acetate (Cat# 106268, Merck). The acetate standard was also used for a spike-in experiment to further confirm the identification.

#### 2.9.3 Batch correction, normalization, and statistical and bioinformatic analyses of metabolomics data

Targeted UHPLC-MS/MS data and ^1^H NMR data were measured within two batches and therefore, they were batch effect corrected using the EigenMS method on the MetaboAnalyst 6.0 platform (https://www.metaboanalyst.ca/). In the case of UHPLC-MS/MS, data collected with negative and positive modes were separately batch-corrected before combining the data. Single missing values within different groups were replaced with 1/5 of the minimum detected values. Metabolite concentrations/abundances were log_2_ transformed for statistical testing with the Student’s t-test with significance set at *p* < 0.05 and fold change > I1.1I.

MetaboAnalyst 6.0 platform was used for the metabolite overrepresentation analysis. Names of the significantly altered metabolites were uploaded. Default settings were used (SMPDB pathway and ‘*Only use metabolites sets containing at least 2 entries*’ option were used).

### 2.10 ROS-experiments and -measurements

ROS levels from the HSkM cells and HUVECs were measured following earlier published protocol (36) with minor modifications. Briefly, 2’,7’-dichlorodihydrofluorescein diacetate probe (DCFH-DA, MecChemExpress, Cat# HY-D0940), was added in the culture medium at a concentration of 10 µM and incubated during the 5-hour mechanical loading or control time. For cells in the antioxidant group, a 100 µM dose of dithiothreitol (DTT) was also added to the medium at the same time as the DCFH-DA. In H_2_O_2_ experiments, the DCFH-DA probe was first incubated 15 minutes without H_2_O_2_, and then together with 250 µM dose of H_2_O_2_ (Merck, Cat# 31642-M) for 30 minutes. Thereafter, cells were rinsed three times with PBS. Then, cells were lysed to 120 µl RIPA buffer (50 mM Tris-HCl, 150 mM NaCl, 1% Triton X-100, 0.1% SDS, 0.5% sodium deoxycholate). Cell lysates were centrifuged at 21,130 x g for 10 min at 4 °C. Thereafter, 85 µl of lysate was transferred to black 96-well plate. Fluorescence intensity was measured using an Agilent BioTek Synergy H1 microplate reader at an excitation wavelength of 485 nm and an emission wavelength of 530 nm.

### 2.11 ^13^C-glucose labeling experiments and analyses

Glucose-free and phenol red-free endothelial cell medium (Cat# H1168GPF, Cell Biologics) medium including the medium kit’s supplements, was supplemented with 5.5 mM U-^13^C_6_-D-glucose (Cat# CLM-1396-1, Cambridge Isotope Laboratories, isotopic purity 99%). This medium was changed to the cells just before the initiation of the experiment. Intracellular metabolites were extracted with the same protocol as for targeted UHPLC-MS/MS metabolomics, but this time two chambers were pooled instead of three. Media metabolites were extracted with the same protocol as for ^1^H NMR metabolomics described above, except now 100 µl of media from each chamber was mixed with 200 µl of methanol. Intracellular and medium metabolites were reconstituted into 45 µl and 400 µl of 50-50 acetonitrile-water solution, respectively. Pooled quality control (QC) samples were prepared by transferring a small aliquot of each sample into a vial (separate QC samples for intracellular and medium metabolites). Selected amino acids were analyzed with the same method as in targeted UHPLC-MS/MS metabolomics described above but the MRM transitions were modified (Supplementary Table S1), allowing analysis of the isotopomers of the selected amino acids. The data was processed and integrated with MRMPROBS_v.3.66 software (37). Phosphoserine and glycine were processed and integrated with Agilent Qualitative Navigator B.08.00 software because of their lower abundance. Isotope natural abundances were corrected with IsoCor 2.2.2 software (38) as earlier reported (39).

### 2.12 PHGDH overexpression

The PHGDH overexpression construct pLJM5-WT PHGDH was a gift from David Sabatini (Addgene plasmid #83901; http://n2t.net/addgene:83901; RRID: Addgene_83901) (40). Lentivirus particles were prepared and distributed by Biomedicum Virus Core facility supported by HiLIFE and the Faculty of Medicine, University of Helsinki, and Biocenter Finland. The control construct (denoted as ‘Control’ in the figure legends) clone SHC002 (Scrambled shRNA in pLKO.1) was from Sigma Aldrich MISSION shRNA library, distributed by Genome Biology Unit core facility supported by HiLIFE and the Faculty of Medicine, University of Helsinki, and Biocenter Finland. The Biomedicum Virus Core facility also determined the p24 capsid protein concentration with an enzyme-linked immunosorbent assay (ELISA). p24 concentrations were used to calculate the number of virus particles needed for transduction (1 pg of p24 = 100 transducing units).

HUVECs were cultured in T75 flasks. After reaching 50% confluence, 10 ml of medium containing lentiviruses (MOI = 50) and polybrene (5 μg/ml), was added to the cells. Polybrene (Cat# TR-1003, Sigma-Aldrich) was used to improve infection efficiency. HUVECs were incubated with lentiviruses for 24 hours. After transduction, the cells were resuspended in a fresh endothelial cell medium, cultured overnight, and used for further experiments. HUVECs expressing scrambled shRNA and HUVECs without virus treatment were used as controls for the PHGDH overexpression construct.

### 2.13 Chemical PHGDH inhibitors

For chemical inhibitor experiments, two different PHGDH inhibitors, NCT-503 (Cat# HY-101966) and CBR-5884 (Cat# HY-100012), were obtained from MedChemExpress. Inhibitor stock solutions were prepared in DMSO, and DMSO was also used as vehicle control. The inhibitors were used in concentrations of 25 or 50 µM, as previously used for different cell types (20, 41–43). Further details of inhibitor experiments are described in the next two sections.

### 2.14 Protein extraction and Western blotting

HUVECs expressing either the PHGDH overexpression system or shControl gene were cultured in 6-well plates (5000 cells/cm^2^) in endothelial cell medium supplemented with 1 µl/ml (vol/vol) ^14^C-[1]-L-phenylalanine (see section 2.7). After 48 hours, cells were homogenized in lysis buffer containing 20 mM HEPES (pH 7.4), 1 mM EDTA, 5 mM EGTA, 10 mM MgCl_2_, 100 mM β-glycerophosphate, 1 mM Na_3_VO_4_, 1 mM DTT, 1% TritonX-100, and supplemented with protease and phosphatase inhibitors (Cat# 78442, Thermo Fisher Scientific). Cell lysates were centrifuged for 5 min at 13,000 x g, 4 ℃, and the supernatant was collected. Separate aliquots of the supernatants were used for western blotting and analysis of ^14^C-[1]-L-phenylalanine incorporation into proteins with liquid scintillation counting (see *Carbon-14 labeling* -section). Western blot analysis of PHGDH was conducted by using low-fluorescence PVDF membranes and anti-PHGDH antibody (1:5000 dilution, Cat# HPA021241, Sigma-Aldrich) as previously (20) with anti-GAPDH (1:10 000 dilution, Cat# ab9485, Abcam) as a loading control.

### 2.15 Statistical testing in non-*omics* analyses

Statistical tests were conducted using GraphPad Prism 10.3.1 (Dotmatics). N-size was mostly ≥6, and normality was tested with Shapiro-Wilk test. In few situations with lower sample sizes (time-course and LDH analyses or when measuring EdU from both sub-confluent and confluent cells), normality was assumed. Student’s t-test or Mann-Whitney U-test was used when two groups were compared depending on the result of Shapiro-Wilk test. One-way ANOVA with Fisher’s least significant difference test was used when more than two groups were compared. The main effects and interaction of two factors were examined using two-way ANOVA. Group difference was considered statistically significant with *p* ≤ 0.05.

## 3 Results

### 3.1 Mechanical loading induced cell type-specific transcriptomic changes mirroring exercise-induced responses

We first investigated the transcriptomic changes in two human endothelial cell types (HUVEC & HCMEC) as well as human skeletal muscle cells (HSkM) (Fig. 1A). The used mechanical loading scheme did not induce detachment of the cells, significant membrane damage, or cell death in either muscle or endothelial cells as determined by total RNA yield and the extracellular lactate dehydrogenase (LDH) activity assay, as well as microscopic images (Supplementary Fig. 1). Differential gene expression analysis demonstrated that all three cell types presented significant changes in their transcriptomes in response to the mechanical loading, as demonstrated by the principal component analysis (PCA) (Fig. 1B). The transcriptomic response was the largest in HUVECs, with the total number of differentially expressed genes being approximately 3-fold larger than in HCMECs or HSkM cells (Fig. 1C). In HUVECs, 717 genes were significantly upregulated, and 779 genes significantly downregulated (log_2_FC ≥ 0.5 & adj. *p* ≤ 0.05), for HCMECs, 241 genes were upregulated, and 261 genes were downregulated, and in HSkM cells 262 genes were upregulated and 126 genes were downregulated. The ten most upregulated and downregulated genes by indexing in order of magnitude and consistency of change (Log_2_FC and adj. *p*) are annotated in Figure 1D for each cell type. For the complete lists of differentially expressed genes and their full names, please refer to the Supplementary Table S2.

**Figure 1.**
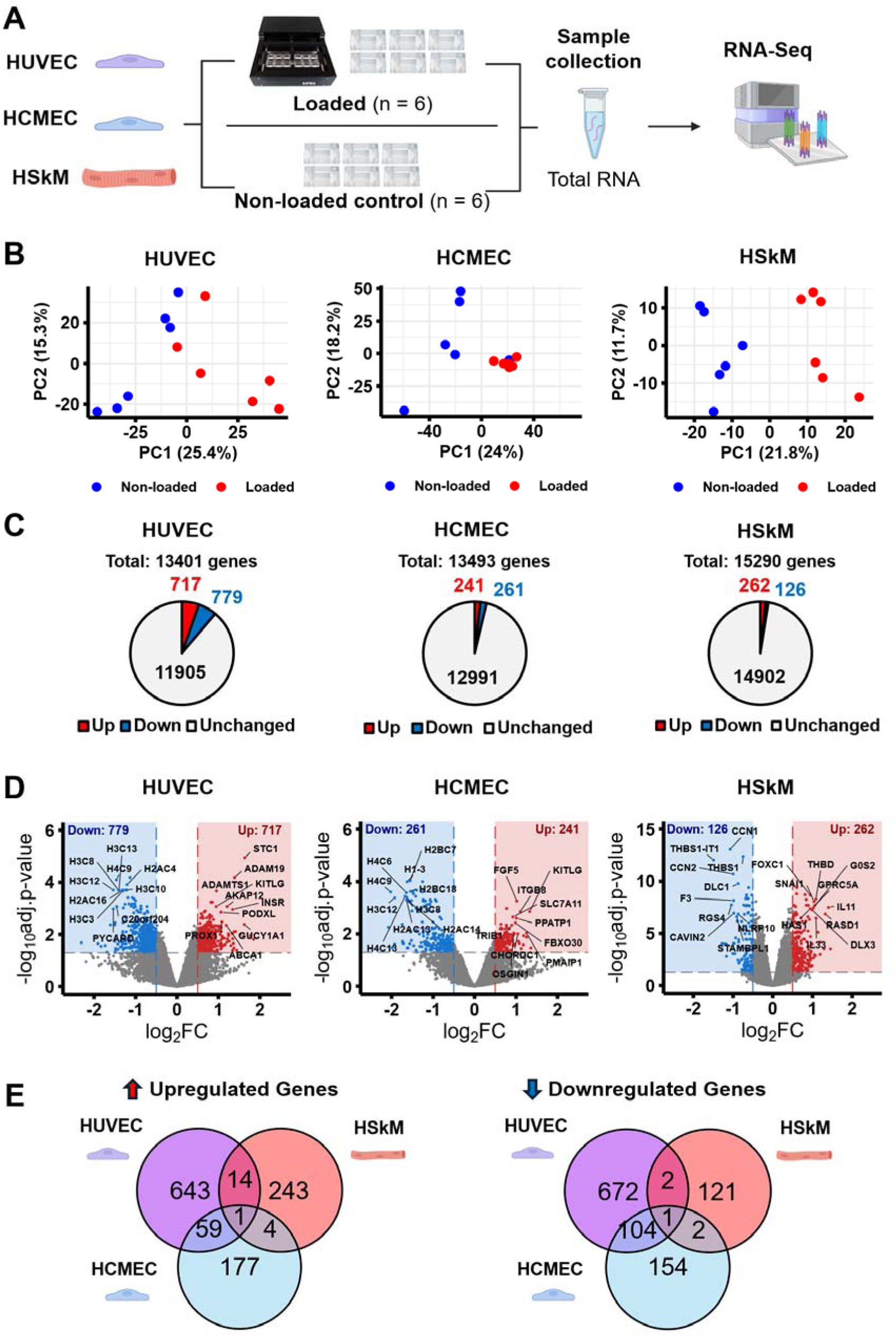
Changes in gene expression of human endothelial cells (HUVECs, HCMECs) and muscle cells (HSkM) in response to mechanical loading. **A)** Schematic representation of experimental design. The experiment was repeated for each cell type separately. Cells from one stretching chamber were used as one sample for RNA-Seq. **B)** PCA plots of samples in each cell type, respectively. **C)** The total number of detected genes and the number of genes that were either up- or downregulated (log_2_FC ≥ 0.5 and adj. *p* ≤ 0.05) for each cell type. **D)** Volcano plots illustrating the amounts of differentially expressed and unchanged genes in all three cell types. Top 10 up- and downregulated genes by magnitude and consistency are annotated **E)** Venn diagrams showing the number of shared up- and downregulated genes in all three cell types.

When comparing overlaps in both up- and downregulated genes in all three cells, we noticed that the two endothelial cells clearly demonstrated similarities in their gene expression, whereas the overlaps between endothelial cells and muscle cells were small (Fig. 1E). Only one gene was upregulated (*CREB5*), and one gene was downregulated (*FAM43A*) in all three cell types in response to mechanical loading. For the complete lists of overlapping differentially expressed genes, please refer to Supplementary Table S3. These results demonstrated greater transcriptomic responses to mechanical loading in endothelial cells than in muscle cells. Large overlaps in gene expression could be seen between the endothelial cell types, whereas endothelial cells and muscle cells responded differently, highlighting the cell type-specificity.

Gene Set Enrichment Analysis (GSEA) results revealed that the two endothelial cell types showed highly similar pathway enrichment after mechanical loading (Supplementary Table S4). Twenty Gene Ontology: Biological Processes (GO:BP) and four Reactome pathways were similarly upregulated in HUVEC and HCMEC, and 27 GO:BP, 83 Reactome and three KEGG pathways were similarly downregulated. The GSEA results of HUVECs showed upregulation of cell maturation and EC junction-related pathways such as *Endothelial cell development, Establishment of endothelial barrier, Cellular response to nitrogen compound* and *Cell junction organization* (Fig. 2A). Out of the GO:BP pathways *Endothelial cell development* and *Establishment of endothelial barrier*, the same 18 genes were significantly upregulated in HUVECs in response to mechanical loading (Fig. 2B). For muscle cells, upregulated pathways were mostly related to oxidative phosphorylation (OXPHOS) and ribosome biogenesis (Fig. 2D). Many of the GSEA findings were also reproduced by the gene set overrepresentation analysis (ORA) (Supplementary Fig. 2).

**Figure 2.**
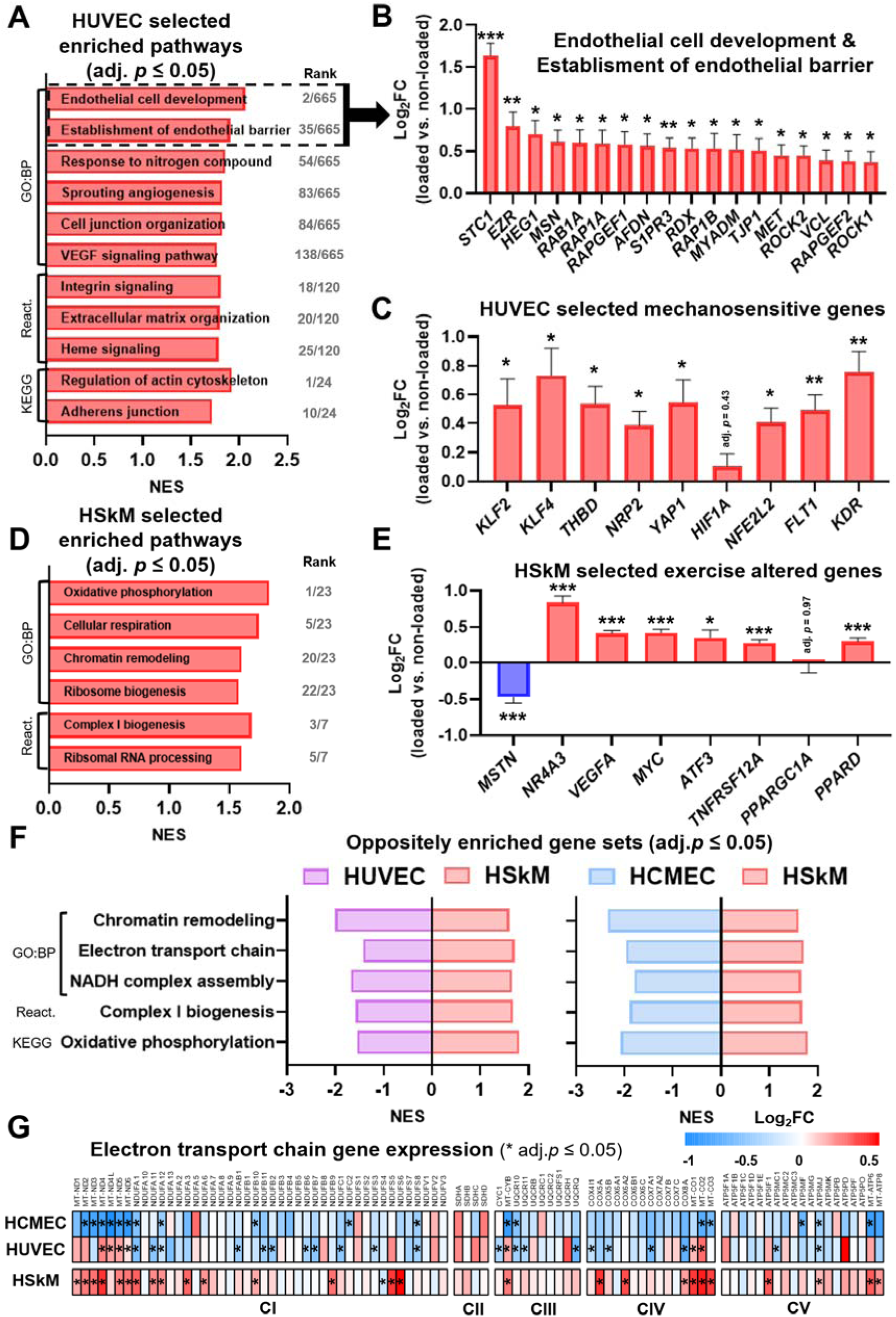
Transcriptomic changes of endothelial and muscle cells in response to mechanical loading. **A)** Selected positively enriched gene sets in HUVECs from GO:BP, Reactome and KEGG database terms. **B)** Upregulated individual genes in HUVECs in GO:BP *Endothelial cell development & Establishment of endothelial barrier* -pathways. **C)** Selected individual mechanosensitive endothelial genes in HUVECs. **D)** Selected positively enriched gene sets in HSkM cells from GO:BP and Reactome database terms. **E)** Selected individual exercise-altered genes in HSkM cells. **F)** Five gene sets from GO:BP, Reactome and KEGG terms, that were significantly downregulated in both endothelial cell types and upregulated in muscle cells. **G)** Heatmap depicting expression of individual genes in all electron transport chain complexes in response to mechanical loading. NES = Normalized enrichment score. GO:BP = Gene ontology:Biological process. Rank = Rank of gene set by NES in all statistically significant (adj. *p* ≤ 0.05) up- or downregulated gene sets in their respective database. Data are presented as mean ± SEM. n = 6 per group in all the analyses. *\*adj*. *p <* 0.05*, **adj. p <* 0.01*, ***adj. p <* 0.001.

To evaluate the external validity of our exercise-on-a-dish model, we analyzed the expression of common exercise-induced and mechanosensitive genes. In endothelial cells, we examined the expression of *KLF2/4*, *THDB*, *NRP2*, *YAP1*, *HIF1A*, and *NFE2L2* and VEGF receptor 1 & 2 encoding genes *FLT1* and *KDR* (Fig. 2C), which have been shown to be induced in ECs in response to flow-induced shear stress (44–46). Indeed, we found that from these genes all but *HIF1A* were significantly upregulated in HUVECs in response to the uniaxial mechanical loading. In HSkM cells, we specifically focused on exercise-altered genes *MSTN*, *NR4A3, VEGFA, MYC, ATF3,* and *TNFRSF12A* (Fig. 2E), and observed similar changes in these genes after mechanical loading as have been shown in response to resistance exercise in vivo (32, 47–49). We additionally looked at the expression of OXPHOS-related genes *PPARGC1A* and its upstream regulator *PPARD* (50–52), and found *PPARD* to be upregulated after mechanical loading, while there was no change in *PPARGC1A* expression (Fig. 2E). Together, these findings suggest that our *in vitro* mechanical loading model mimics exercise and flow-induced mechanical stress in both endothelial and muscle cells.

When comparing gene set enrichment of the endothelial cell types to muscle cells, we found multiple pathways enriched in opposite directions in response to mechanical loading (Supplementary Table S5). In HUVECs and HSkM cells 12 GO:BP, three Reactome and two KEGG pathways were oppositely enriched and the same was true for HCMECs and HSkM cells in 14 GO:BP, four Reactome and three KEGG pathways. Five of these pathways were found to be oppositely regulated when comparing both endothelial cell types to muscle cells. These pathways were related to cellular respiration, metabolism, and chromatin remodeling (Fig. 2F). Most prominently, OXPHOS-related pathways were downregulated in endothelial cells, while upregulated in muscle cells. Closer inspection revealed that gene expression was specifically affected in the same fashion throughout the electron transport chain (ETC)-complexes I-V (Fig. 2G).

Together, in endothelial cells mechanical loading induced positive enrichment of pathways related to maturation and barrier establishment as well as upregulation of known mechanosensitive genes. In muscle cells mechanical loading upregulated OXPHOS and ribosome biogenesis pathways, as well as changes in typical resistance exercise-regulated genes, suggesting that mechanical loading alone recapitulates several common exercise effects seen *in vivo*. The transcriptomic results clearly demonstrated cell type-specific responses induced by mechanical loading. This was seen especially in the opposite expression of ETC-complex genes, which were downregulated in endothelial cells and upregulated in muscle cells.

### 3.2 Mechanical loading blunts proliferation and increases junctional stability of endothelial cells

Based on the transcriptomic results, we were interested in seeing whether the establishment of the endothelial barrier would be present at the structural level. Indeed, we found that mechanical loading led to the re-arrangement of endothelial cells perpendicular to the direction of stretch and increased the amount of cell-cell contacts (Fig. 3A). It also increased intensity of endothelial junction proteins VE-Cadherin and Claudin 5 and similarly, but somewhat less clearly also Tight Junction Protein 1 (ZO-1) (Fig. 3A). Mechanical loading decreased proliferation of HUVECs, analyzed with both flow cytometry- and microscopy-based assays (Fig. 3B-C), whereas overall protein synthesis was not affected (Fig. 3D). These findings, together with the transcriptomic results (Fig. 2B), suggest that mechanical loading aligns the cells, induces junctional stability and attenuates proliferation of endothelial cells, directing the cells to a more quiescent phenotype. To translate our results to an *in vivo* situation, we applied the MetaMEx database (32) to examine whether the *Endothelial Cell Development* -pathway is induced in response to exercise. Indeed, we found several of these genes upregulated in human skeletal muscle after acute aerobic exercise (Supplementary Fig. 3).

**Figure 3.**
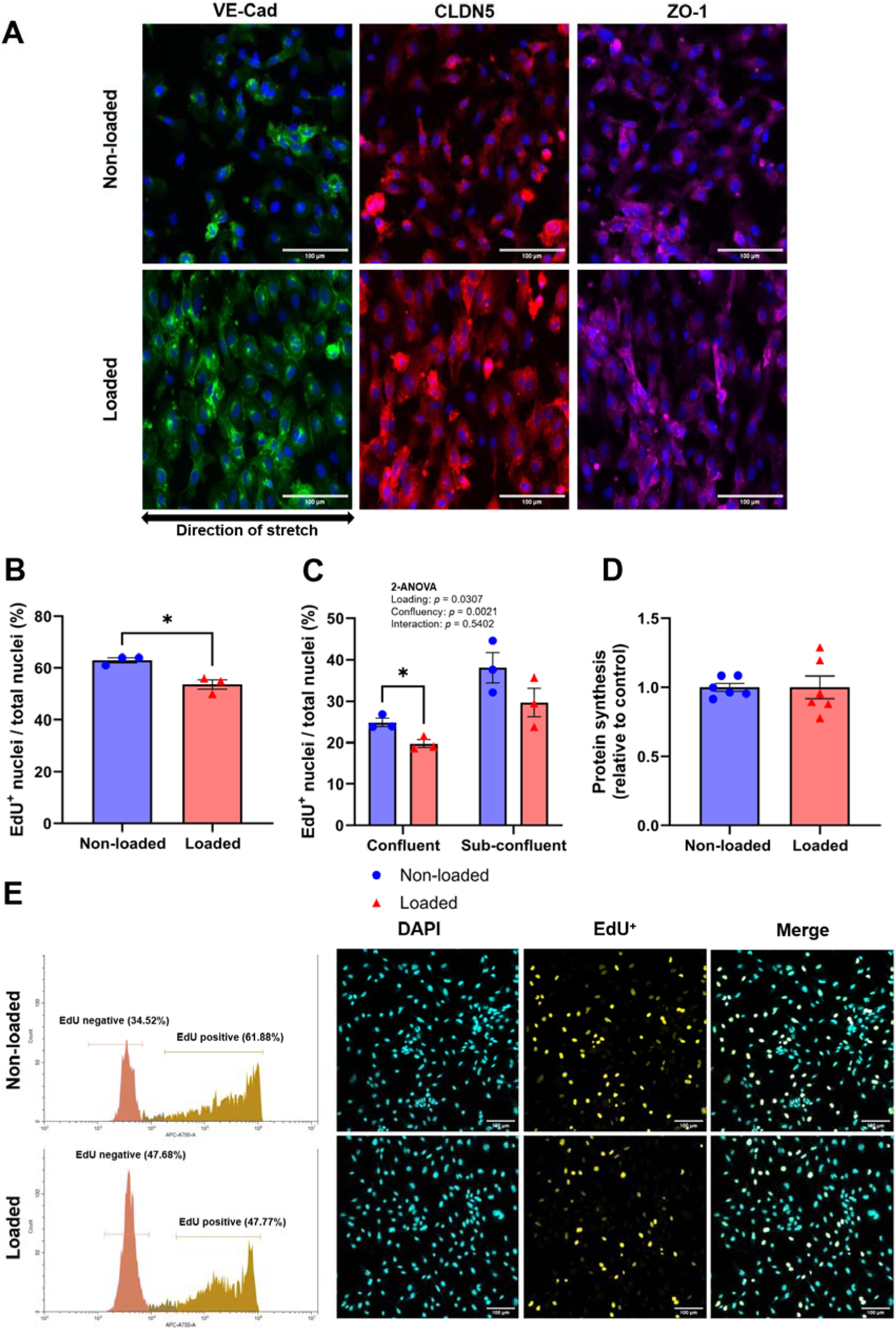
Effects of mechanical loading on endothelial cell (HUVEC) proliferation, protein synthesis and morphology. **A)** Confocal microscopy images showing the effect of mechanical loading on cell morphology in endothelial cells stained for junctional proteins VE-Cadherin (VE-Cad), Claudin 5 (CLDN5) and Tight Junction Protein 1 (ZO-1 / *TJP1*). **B)** EdU-positive cells analyzed with flow cytometry (n = 3). **C)** EdU-positive cells in confluent and sub-confluent cell densities were analyzed with confocal microscopy (n = 3). **D)** Protein synthesis assessed by relative ^14^C-phenylalanine incorporation (n = 6). **E)** Representative flow cytometry histograms showing EdU negative and positive cell populations, and next to them are representative confocal microscopy images showing all nuclei (DAPI) and EdU positive nuclei. Data are presented as mean ± SEM, and individual values are plotted when applicable. \**p <* 0.05 (Student’s t-test).

3.3 Both cell types released acetate in response to mechanical loading

Next, we compared extracellular (media) metabolite changes in HUVECs and HSkM cells in response to mechanical loading. HCMECs were excluded because these analyses required more sample material and resources. Also, based on the transcriptomic results, HUVECs were a more promising candidate for further investigation as their transcriptomic changes more closely mirrored what has been observed in vivo (18). For this, culture media metabolites were analyzed with fully quantitative ^1^H NMR spectroscopy. From the extracellular metabolites (conditioned media), 3/29 were altered in HSkM cells, and 2/34 in HUVECs (Fig. 4A and Supplementary Table S6).

**Figure 4.**
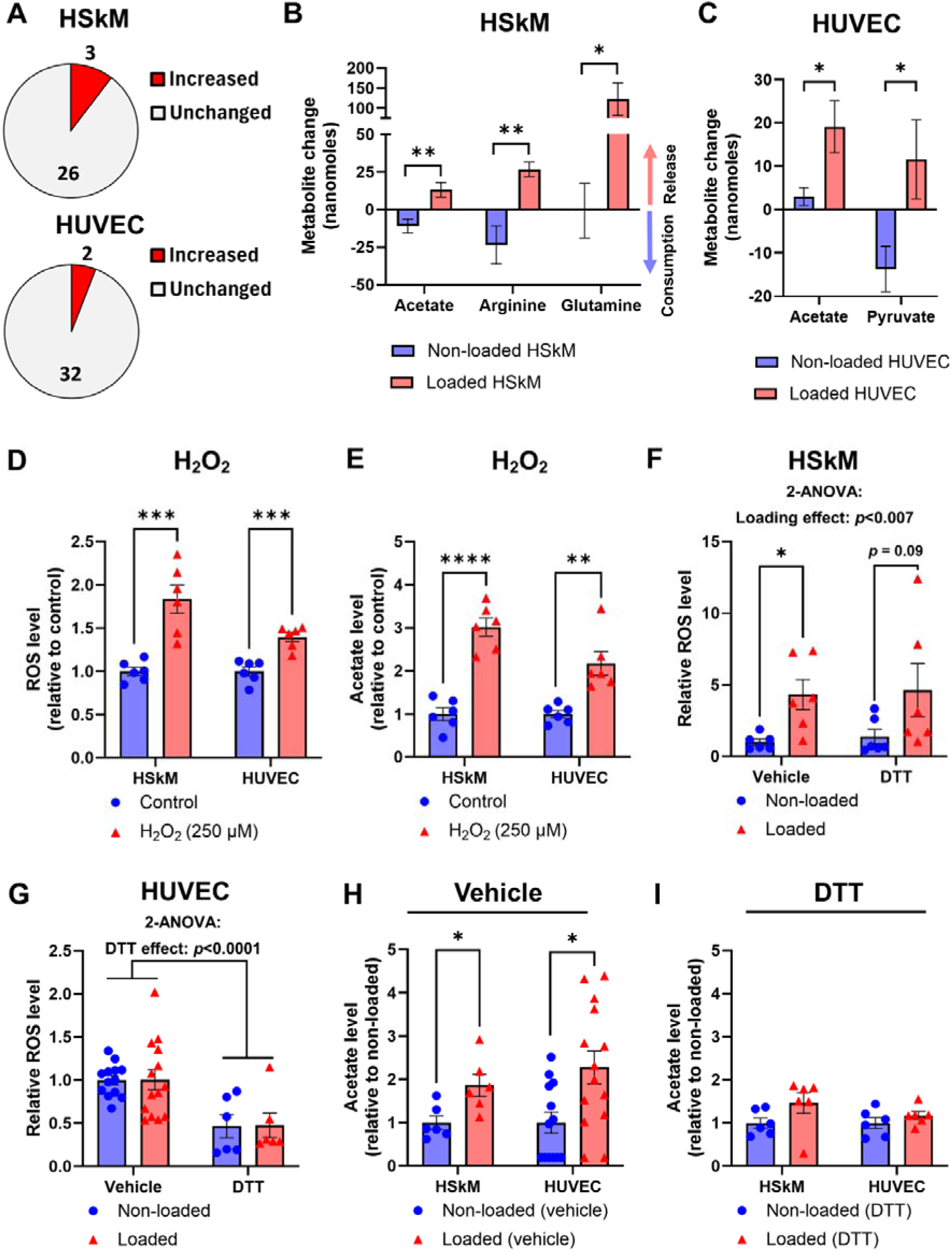
Analysis of extracellular (media) metabolites and intracellular reactive oxygen species (ROS). **A)** The number of significantly (*p*<0.05 and FC>I1.1I) changed and unchanged media metabolites in response to mechanical loading measured with ^1^H NMR spectroscopy. **B-C)** Significantly released metabolites (nanomoles per 4 cm^2^ of cells over 5 hours) by HSkM cells or HUVECs in response to mechanical loading. Analysis of both medium incubated with cells and medium incubated without cells enabled estimation of metabolite release or consumption. **D)** Analysis of intracellular ROS in response to H_2_O_2_ treatment. **E)** Analysis of media acetate levels in response to H_2_O_2_ treatment measured with UHPLC-MS. **F-G)** ROS levels in response to mechanical loading and an antioxidant, dithiothreitol (DTT), in HSkM cells and HUVECs, respectively. **H)** Acetate levels in culture media in response to mechanical loading measured with UHPLC-MS. **I)** DTT suppressed acetate release to culture media in response to loading. n = 6-14 depending on group and analysis. \**p < 0.05, **p < 0.01, ***p < 0.001,* **** *p* < 0.0001. Student’s t-test was used except in Panels E and H for HUVEC comparisons and Panel I for HSkM comparison; the Mann-Whitney U test was used because of a violation of normality.

Next, we calculated net metabolite release or consumption by considering the metabolite concentrations in media incubated without cells (i.e., cell-free media). We found that both HUVECs and HSkM cells released acetate into the media in response to mechanical loading (Fig. 4B-C). One potential source of acetate after exercise and mechanical loading could be its formation from pyruvate by ROS (53), since ROS production is induced in muscle by exercise and by stretching of endothelial cells and skeletal muscle ex vivo (54–56). Here, we tested whether our mechanical loading approach induces acetate release in a ROS-dependent manner. First, we validated that addition of H_2_O_2_ increased measured ROS levels in both cell types (Fig. 4D). Moreover, H_2_O_2_ treatment alone increased acetate levels in culture media (Fig. 4E), supporting the ROS-mediated mechanism also in the loading situation. Next, we found that mechanical loading induced an elevation in ROS levels in HSkM cells but not in HUVECs (Fig 4F-G). However, an antioxidant treatment (DTT) prevented mechanical loading-induced acetate secretion in both cell types (Fig. 4H-I). Together, these findings suggest that although cell-type differences were observed, the acetate release in response to mechanical loading may be at least partially ROS-dependent in both cell types.

### 3.4 Intracellular metabolite changes were greater in endothelial cells than in muscle cells

We analyzed a targeted list of intracellular metabolites from HSkM cells and HUVECs using UHPLC-MS/MS (Fig. 5A). In HSkM cells and HUVECs, 3/35 and 11/38 intracellular metabolites were altered in response to mechanical loading, respectively (Fig. 5B). The intracellular metabolite alterations were thus greater in HUVECs than in HSkM cells, which is in line with the transcriptome results. In muscle cells, intracellular levels of glucose 6-phosphate and phosphocreatine decreased while alanine levels increased in response to mechanical loading (Fig. 5C). In HUVECs, intracellular levels of phosphoenolpyruvate, fructose 6-phosphate, and α-ketoglutarate decreased while phosphoserine, serine, asparagine, threonine, proline, alanine, acetyl-CoA, and tryptophan levels increased (Fig. 5D). The two mostly increased metabolites were phosphoserine and serine (Fig. 5D), whereas the serine synthesis pathway gene expressions were unaffected (Supplementary Table S2). Proline and α-ketoglutarate were the two most consistently (smallest *p-*values) altered metabolites. The observed increase in proline and decrease in α-ketoglutarate may indicate that proline was synthesized using α-ketoglutarate.

**Figure 5.**
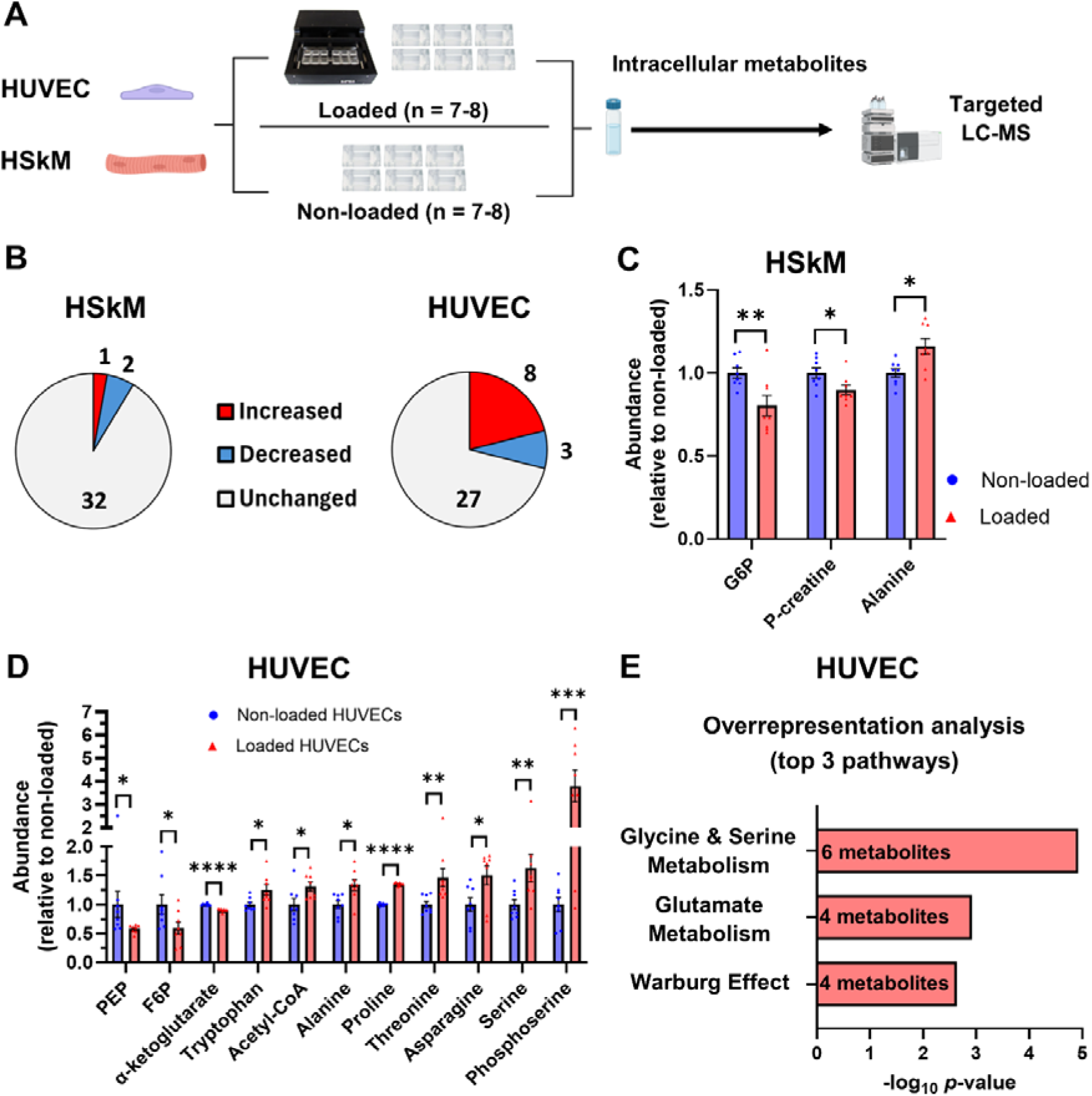
Intracellular metabolite changes in human skeletal muscle cells (HSkM) and HUVECs in response to mechanical loading**. A)** Intracellular metabolites were analyzed using the targeted UHPLC-MS/MS method. Cells from three stretching chambers were pooled for one sample (n = 7-8 per group). **B)** The number of significantly (*p*<0.05 and FC>I1.1I) increased, decreased, and unchanged metabolites of HSkM cells and HUVECs. **C)** Significantly changed intracellular metabolites in HSkM cells. **D)** Significantly changed intracellular metabolites in HUVECs. **E)** Metabolite overrepresentation analysis using the significantly altered intracellular metabolites of HUVECs. In panel E, the number of metabolites indicates how many significantly altered metabolites per pathway were detected. Data are presented as mean ± SEM, and individual values are plotted. P-creatine = Phosphocreatine, G6P = Glucose 6-phosphate, PEP = Phosphoenolpyruvate, F6P = Fructose 6-phosphate. \**p <* 0.05*, **p <* 0.01, *\*\*\*p* < 0.001, **** p < 0.0001 (Student’s t-test).

Next, we conducted a metabolite overrepresentation analysis using the significantly altered intracellular metabolites. The overrepresentation analysis of HSkM cells did not yield any significant results because of the low number of altered metabolites. In HUVECs, the most enriched pathways were *Glycine and serine metabolism*, *Glutamate metabolism*, and the *Warburg effect* (Fig. 5E).

### 3.5 Mechanical loading increased the synthesis of serine, alanine, and proline from glucose in endothelial cells

As serine metabolism was the most highlighted pathway in metabolomics analysis of mechanically loaded HUVECs, and its role in survival and growth has been demonstrated (57), we next conducted dynamic labeling experiments with ^13^C-(U)-glucose (Fig. 6A) to assess whether the serine synthesis rate is affected by mechanical loading. We found that there were significantly more total (unlabeled + labeled) and newly synthesized serine and phosphoserine at multiple time points after mechanical loading (Fig. 6B-E). This indicates that the serine synthesis was indeed increased in response to mechanical loading in HUVECs. Total serine levels did not change in the media (Fig. 6F), whereas mechanically loaded HUVECs released significantly more labeled serine into the culture media (Fig. 6G), supporting the interpretation that serine synthesis was increased by mechanical loading.

**Figure 6.**
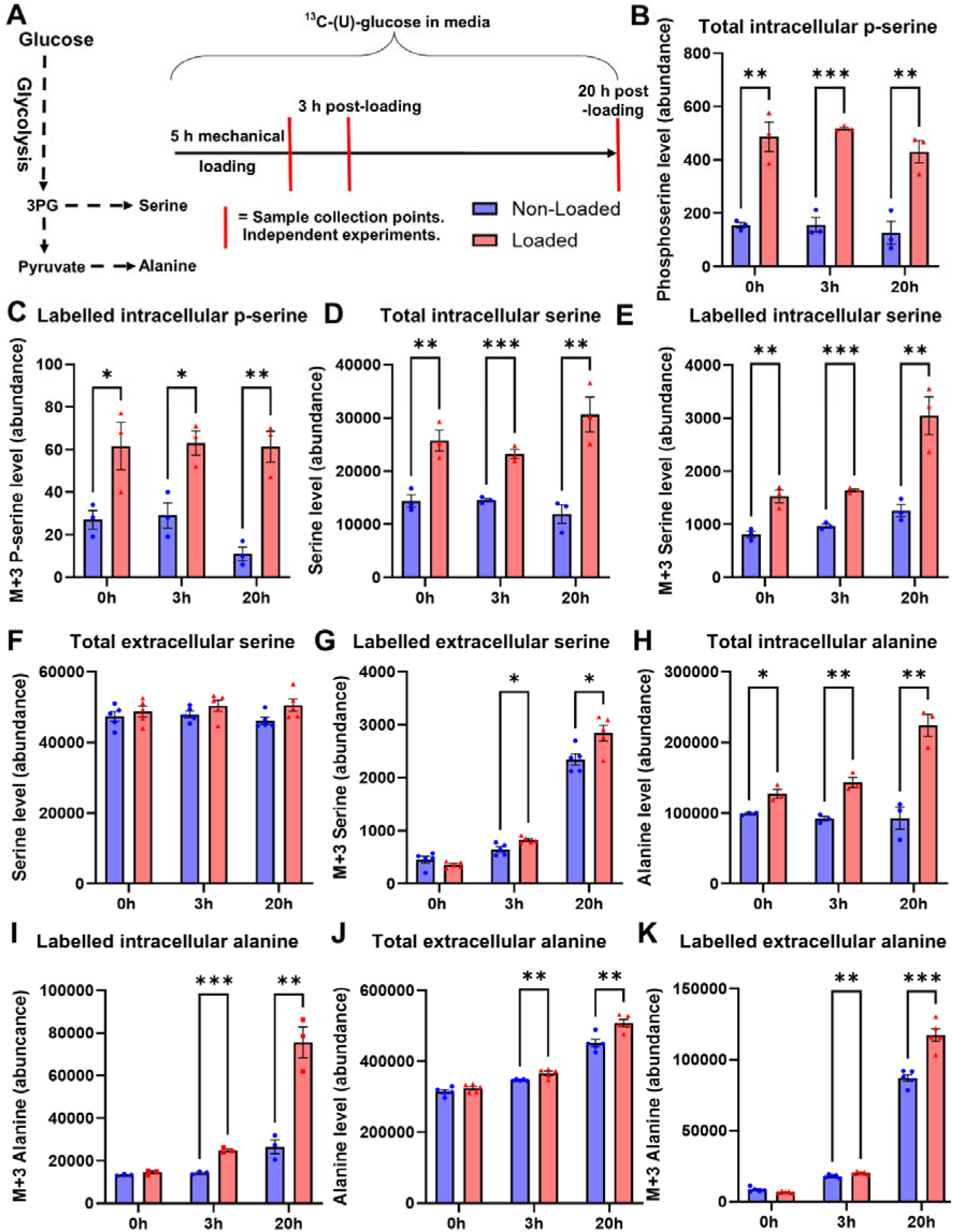
Mechanical loading increased glucose-derived carbon incorporation into serine and alanine in HUVECs. **A)** A schematic of the ^13^C-(U)-glucose experiment. **B)** Total intracellular serine level (unlabeled + labeled). **C)** Newly synthesized (M+3) serine level. **D)** Total intracellular phosphoserine (P-serine) level (unlabeled + labeled). **E)** Newly synthesized (M+3) phosphoserine level. **F)** Total media (extracellular) serine level (unlabeled + labeled). **G)** Newly synthesized (M+3) media serine level. **H)** Total intracellular alanine level (unlabeled + labeled). **I)** Newly synthesized (M+3) alanine level. **J)** Total media (extracellular) alanine level (unlabeled + labeled). **K)** Newly synthesized (M+3) media alanine level. Sample sizes were n = 3 per group per time point in intracellular analyses and n = 5 in media analyses. Individual values are plotted, and data are presented as mean ± SEM. \**p <* 0.05, *\*\*p <* 0.01, *\*\*\*p* < 0.001 (Student’s t-test).

We also assessed the synthesis of alanine, proline, glycine, aspartate, asparagine, glutamine, glutamate, and arginine. We observed higher levels of labeled and total alanine in loaded HUVECs and their media (Figure 6H-K). Similar results were also observed for proline (Supplementary Fig. 4A-C). These data indicate increased alanine and proline synthesis from glucose, along with serine synthesis, in response to mechanical loading.

There was significantly less labeled glycine in loaded HUVECs at 3 h and 20 h time points and total glycine at the 0 h time point (Supplementary Fig. 4D-E), but no differences in labeled or total glycine release into media (Supplementary Fig. 4F-G) compared to non-loaded controls. We did not find significant differences in labeled aspartate and glutamate (Supplementary Fig. 5). In the case of asparagine, arginine, and glutamine, ^13^C-labeled molecules were not detected after isotope natural abundance correction, suggesting that HUVECs did not use glucose to produce these amino acids in these nutritional conditions (Supplementary Table S7). Overall, the results suggest that mechanical loading induces the synthesis of serine, alanine, and proline in endothelial cells.

### 3.6 Serine synthesis pathway enzyme phosphoglycerate dehydrogenase (PHGDH) is critical for endothelial cell anabolism

Finally, as we found that mechanical loading increased serine synthesis, we wanted to extend the previous findings on the role of the serine synthesis pathway in endothelial cells (57). For this we used a lentiviral vector to overexpress PHGDH, the rate-limiting enzyme of the serine synthesis pathway (Supplementary Fig. 6A). Overexpression of PHGDH led to a ∼10% increase in protein synthesis (*p* < 0.01, Supplementary Fig. 6B) and a trend for increased proliferation (*p* = 0.088, Supplementary Fig. 6C) in HUVECs. Next, we used chemical PHGDH inhibitors and evaluated their effects on protein synthesis and proliferation. We discovered that PHGDH inhibitors decreased the protein synthesis and proliferation of HUVECs (Supplementary Fig. 6D-E) without inducing cell death (Supplementary Fig. 6F). We also found that the PHGDH inhibitors decreased glucose-derived carbon incorporation into proteins, RNA, and lipids (Supplementary Fig. 6G-I). Overall, these data suggest that the serine synthesis pathway enzyme PHGDH regulates endothelial cell growth, similarly to what we recently showed in muscle cells (20, 34).

## 4 Discussion

In this study, we unraveled the effects of mechanical loading on skeletal muscle and endothelial cells and identified both shared and cell type-specific responses. Notably, both muscle and endothelial cells significantly increased acetate release into the media and induced gene expression changes typically observed after exercise. However, transcriptomic responses exhibited several contrasting patterns in endothelial and muscle cells. Most strikingly, OXPHOS and electron transport chain (ETC) genes were upregulated in muscle cells but downregulated in endothelial cells. In response to mechanical loading, endothelial cells showed indications of increased junctional stability and maturation. Furthermore, endothelial cells demonstrated more pronounced metabolic responses, particularly in amino acid metabolism. Using a ^13^C-(U)-glucose tracer, we demonstrated that mechanical loading enhanced glucose-derived serine synthesis in endothelial cells. Overexpression of the serine synthesis enzyme PHGDH promoted protein synthesis of endothelial cells, whereas PHGDH inhibitors decreased proliferation and anabolic activity, highlighting the role of de novo serine synthesis from glucose in endothelial cell growth.

Both muscle and endothelial cells released acetate into the media in response to mechanical loading, which was observed with two different analysis methods in multiple independent experiments. Previously, we found that electrically evoked myotube contractions also induced acetate release from murine myotubes (33). In humans, it has been found that acetate is released into the bloodstream (58) and urine (59) after exercise. Moreover, serum acetate concentrations have been found to be greater in physically active individuals compared to inactive individuals (60). Therefore, acetate could act as a possible exerkine (i.e. a signaling molecule released in response to exercise) released by not only muscle cells but also endothelial cells. A potential source of acetate after exercise and mechanical loading could be its formation from pyruvate by ROS (53), since ROS production is induced in muscle by exercise and stretching of endothelial cells and skeletal muscle ex vivo (54–56). Here, we found that an antioxidant (DTT) blunted stretch-induced acetate release in both cell types, suggesting that acetate release may be in part ROS-dependent in both cell types. Interestingly, we could only detect increased ROS in response to mechanical loading in muscle cells, but not in endothelial cells. However, it is possible that mechanical loading induced localized ROS production also in endothelial cells, but this was not observable from total cell lysate. Furthermore, quiescent endothelial cells have also been shown to have improved ROS-scavenging capabilities compared to proliferating endothelial cells (61). Additional potential sources of acetate after mechanical loading could be its formation from acetyl-CoA by certain acyl-CoA thioesterases (62), and by deacetylation of proteins (63), thus it would be interesting to further dissect the physiological role of acetate release. One of the most relevant known functions of acetate is acting as an immunomodulator and promoter of T-cell effector function and regulating differentiation of CD4^+^ T-cell helper cells (64–66). Therefore, acetate release in response to exercise could potentially partly explain the previously observed effects of exercise in enhancing immune system function and increasing CD4^+^ T-helper cell count in humans (67). This would be an interesting research line for the future, because factors secreted by CD4^+^ T-cells appear to be important in angiogenesis, myogenesis, and muscle regeneration (68, 69).

Endothelial cells and muscle cells also demonstrated different and even opposite transcriptomic responses to mechanical loading, most prominently in OXPHOS-related pathways, and specifically in ETC complex genes. In endothelial cells the expression of these genes was downregulated, and it has previously been shown that they primarily rely on glycolysis for ATP production despite having access to oxygen, and that vessel sprouting *in vitro* is regulated by glycolysis (70). OXPHOS-related genes were on the other hand upregulated in muscle cells, and skeletal muscle is known to upregulate mitochondrial biogenesis and oxidative respiration in response to physiological stress, such as exercise (19). One of the well-known mechanisms regulating these exercise adaptations is the AMPK-PGC1α pathway (50). In our study, although PGC1α gene expression was unchanged in muscle cells, we did observe an increase in *PPARD* expression after mechanical loading. PPARD is upstream of PGC1α (51) and has been linked to enhanced oxidative metabolism in skeletal muscle (52). Furthermore, muscle cells also demonstrated an upregulation of the *NR4A3, ATF3, and TNFRSF12A*, which have been shown to belong to the five most upregulated genes in response to resistance exercise (32). For example, *NR4A3* is essential in glucose metabolism and anabolism, such as ribosome biogenesis in primary muscle cells (47). In line with the upregulation of *NR4A3*, we also observed increases in the ribosome biogenesis pathway after mechanical loading, which is a known response to resistance training (71). Moreover, we saw a decrease in myostatin (*MSTN*) expression in muscle cells, which is a common response after a single bout of resistance exercise (48, 49). These results demonstrate that mechanical loading of human skeletal muscle cells recapitulates several of the common effects of exercise seen *in vivo*.

In endothelial cells, transcriptomic results showed that mechanical loading induced upregulation of genes related to endothelial cell development, migration, cell-cell junctions, and endothelial barrier integrity. In comparison to datasets from human exercise studies, many of the endothelial cell development genes were also found to be similarly increased in response to acute aerobic exercise in human skeletal muscle tissue, which is known to be highly vascularized (1, 2, 32). Related to barrier integrity, we also observed that the mechanical loading induced increased cell-cell contacts in endothelial cells, with a notable change in junctional protein detection via immunofluorescence imaging, especially in VE-Cadherin. Moreover, the proliferation rate of endothelial cells was blunted with mechanical loading compared to non-loaded cells. Furthermore, our results showed upregulation of mechanosensitive genes commonly associated with an atheroprotective endothelial phenotype, which are typically induced by flow-mediated mechanical stress (44–46), but here observed after uniaxial cyclic stretching, which could be comparable to skeletal muscle movement. These results suggest that mechanical loading induces endothelial cell quiescence and improves barrier function and integrity. We also found that genes related to VEGF response, including VEGF receptors 1 & 2 (*FLT1 & KDR*), were upregulated in endothelial cells. Recently, a study investigating acute endothelial-specific transcriptional responses in mouse skeletal muscle with bulk RNA-seq of endothelial cell fractions, also found that exercise induced activation of pathways related to endothelial cell development, differentiation, and migration, as well as VEGF response (18). This aligns well with our results and demonstrates the similarities between endothelial cell responses to mechanical loading and exercise, suggesting that mechanical loading of endothelial cells can potentially recapitulate the effects of exercise *in vivo*.

In endothelial cells, we observed upregulation of heme signaling- and endothelial cell maturation-related genes, while serine synthesis increased after mechanical loading. Mechanistically, the overexpression of the serine-synthesizing enzyme PHGDH increased protein synthesis, while commonly used non-competitive PHGDH inhibitors decreased proliferation, protein synthesis, and glucose utilization for endothelial cell anabolism. This is similar to what we recently demonstrated in muscle cells (20, 34). Furthermore, loss of PHGDH has previously been shown to cause lethal defects in angiogenesis in mice, and knockdown of PHGDH impairs endothelial cell proliferation (57). Moreover, PHGDH has been highlighted as essential for the regulation of heme production in endothelial cells (57), and hemin (oxidized heme) supplementation promotes differentiation of endothelial progenitor cells (72). In future studies, the potential link between the serine synthesis pathway, heme signaling, and endothelial cell maturation should be investigated.

### Strengths and limitations

We used two different human endothelial cell types and primary human skeletal muscle cells. The loading induced several common exercise-induced and mechanosensitive genes, suggesting an external validity of the exercise-on-a-dish model. Moreover, several of the transcriptomic responses to mechanical loading were similar between the two endothelial cell types (HUVEC and HCMEC), suggesting that many of the changes can be applied to endothelial cells in general. We also replicated the main metabolite and proliferation results with different methods. For example, acetate release was observed with both ^1^H NMR spectroscopy and UHPLC-MS, and the decrease in proliferation was determined with microscopy- and flow-cytometry-based analyses. In addition, the increased serine, alanine, and proline content and biosynthesis were detected by targeted metabolomics and isotopic tracing from multiple time points.

The study also has limitations. We conducted experiments with monolayers of cells (i.e., two-dimensionally), and therefore, myotubes were not aligned unidirectionally while the mechanical loading was unidirectional. In contrast, smaller endothelial cells could change their alignment against the direction of stretch. This difference in the ability to change alignment may in part explain why we detected fewer molecular changes in myotubes compared to endothelial cells, although myotubes also recapitulated several exercise responses. Overall, the data suggests that the mechanical loading model for two-dimensionally cultured cells is especially suitable for endothelial cells, which also grow in monolayers *in vivo*, while myotubes may need to be loaded three-dimensionally to sufficiently mimic exercise in future studies. The potential importance of the mechanical loading and serine synthesis pathway for improved vascular function should be studied *in vivo* in further studies.

## 5. Conclusion

Mechanical loading induces both common and highly cell type-specific transcriptomic and metabolomic responses in endothelial and skeletal muscle cells. In general, several of the effects recapitulated what is observed *in vivo* after exercise. Both endothelial and muscle cells released acetate in response to mechanical loading, possibly at least in part via ROS-dependent mechanisms, and therefore acetate’s potential role as an exerkine could be an interesting topic for future studies. Mechanical loading induces a shift towards a quiescent endothelial phenotype with decreased proliferation, increased cell-cell contacts and increased expression of genes related to barrier integrity. The metabolic responses to mechanical loading were larger in endothelial cells, revealing increased serine synthesis and upregulation of endothelial cell development as one of the main responses. These results highlight that mechanical stimulus, endothelial cell quiescence, and serine synthesis may be interconnected.

## Author Contributions

**Sakari Mäntyselkä:** Conceptualization, Methodology, Validation, Formal analysis, Investigation, Resources, Writing – Original Draft, Writing – Review & Editing, Visualization. **Erik Niem**i: Conceptualization, Methodology, Validation, Formal analysis, Investigation, Data Curation, Writing – Original Draft, Writing – Review & Editing, Visualization. **Laura Ylä-Outinen:** Conceptualization, Methodology, Investigation, Writing – Review & Editing**. Kalle Kolari:** Investigation, Writing – Review & Editing. **Liina Uusitalo-Kylmälä:** Formal analysis, Investigation, Writing – Review & Editing, Visualization. **Alfredo Ortega-Alonso:** Formal Analysis, Data Curation, Writing – Review & Editing. **Roosa-Maria Liimatainen:** Validation, Investigation, Review of the manuscript. **Vasco Fachada:** Software, Writing – Review & Editing. **Perttu Permi:** Investigation, Writing – Review & Editing. **Elina Kalenius:** Methodology, Writing – Review & Editing, Supervision. **Juha J Hulmi:** Conceptualization, Resources, Writing – Original Draft, Writing – Review & Editing, Supervision, Project administration. **Riikka Kivelä:** Conceptualization, Resources, Writing – Original Draft, Writing – Review & Editing, Supervision, Project administration, Funding acquisition.

## Acknowledgments

We thank Associate Professor Thomas M. O’Connell for valuable advice regarding ^13^C isotopic tracing and for providing the R script for ranking the significantly altered genes. We also thank all the laboratory technicians who helped us during this study. Figures 1A and 5A were created with the licensed version of https://www.biorender.com/.

## Funding

This work was supported by the Research Council of Finland (R.K., Grant number: 348636), the Sigrid Jusélius Foundation (R.K.), the Finnish Foundation for Cardiovascular Research (R.K., L. U-K.), the Emil Aaltonen Foundation (S.M., E.N.), the Finnish Cultural Foundation (K.K. and J.J.H), and the Orion Research Foundation (S.M.).

## Conflicts of Interest statement

The authors declare that they have no known competing financial interests or personal relationships that could have appeared to influence the work reported in this paper.

## Data availability Statement

Filtered non-aligned RNA sequencing data and non-normalized quantified reads are available in the Gene Expression Omnibus (GEO) data repository (accession number: GSE288064). Processed transcriptomics data are shared as Supplementary Excel spreadsheets. Numerical metabolomics and isotopic tracing data are shared as Supplementary Excel spreadsheets (see *Supporting Information*).

## Supporting Information

A PDF file containing Supplementary Figures 1-6. Excel spreadsheets:

- Table S1: MRM-transitions used in ^13^C-(U)-glucose tracing
- Table S2: List of DEGs
- Table S3: Overlapping DEGs
- Table S4: GSEA results
- Table S5: Overlapping GSEA pathways
- Table S6: Metabolomics data
- Table S7: ^13^C-(U)-glucose labeling data

## Notes

### Competing Interest Statement

The authors have declared no competing interest.

### Summary of Updates

Figure 3 was missing from the previous version. It has now been added.

