## Supplementary Figures for "Mechanical loading induces distinct and shared responses in endothelial and muscle cells and reveals exercise-like molecular profiles"

This file includes:

- Supplementary Figure 1
- Supplementary Figure 2
- Supplementary Figure 3
- Supplementary Figure 4
- Supplementary Figure 5
- Supplementary Figure 6

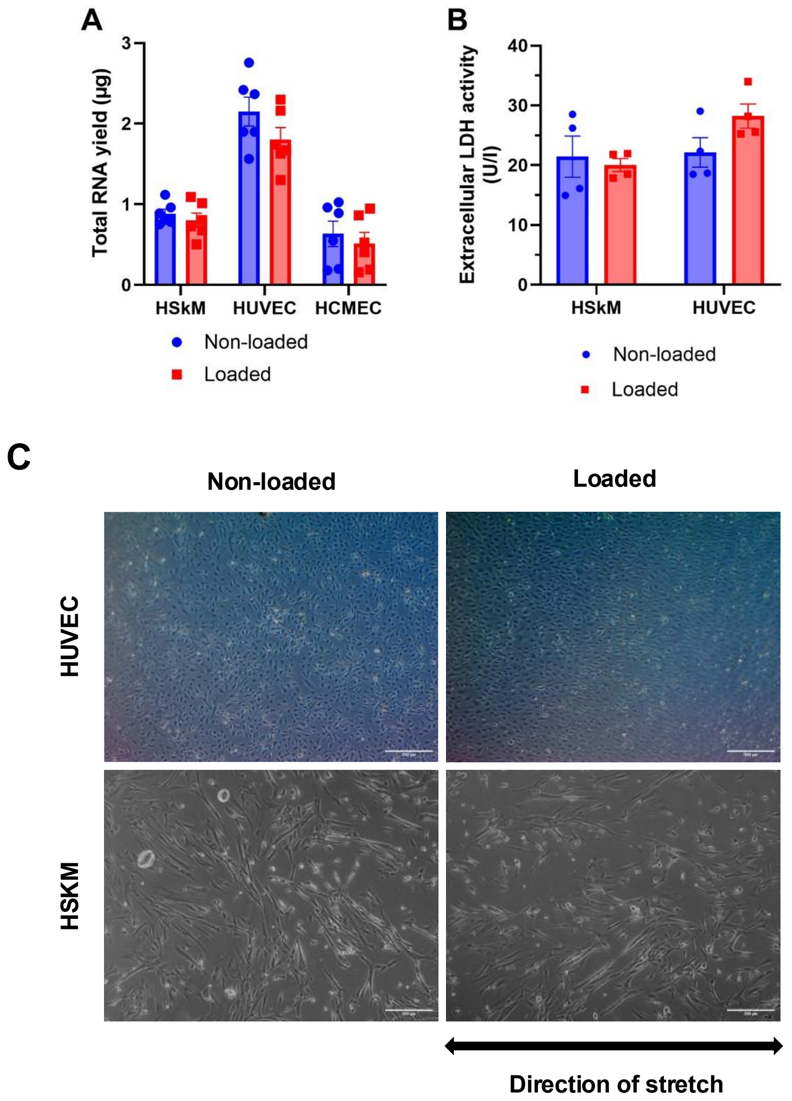


**Supplementary Fig. 1.** Mechanical loading did not induce detachment of the cells, cell, membrane damage, or cell death. **A)** Total RNA yield was similar between the non-loaded and loaded cells, indicating that no significant detachment of cells occurred (n = 6). **B)** Mechanical loading did not significantly affect the extracellular lactate dehydrogenase (LDH) activity, a marker of cell membrane damage and cell death (n = 4). LDH activity was not determined from the media of HCMECs because the media were not collected from those experiments. **C)** Representative light-microscope images of HUVEC and Human skeletal muscle (HSKM) cells before and after mechanical loading.

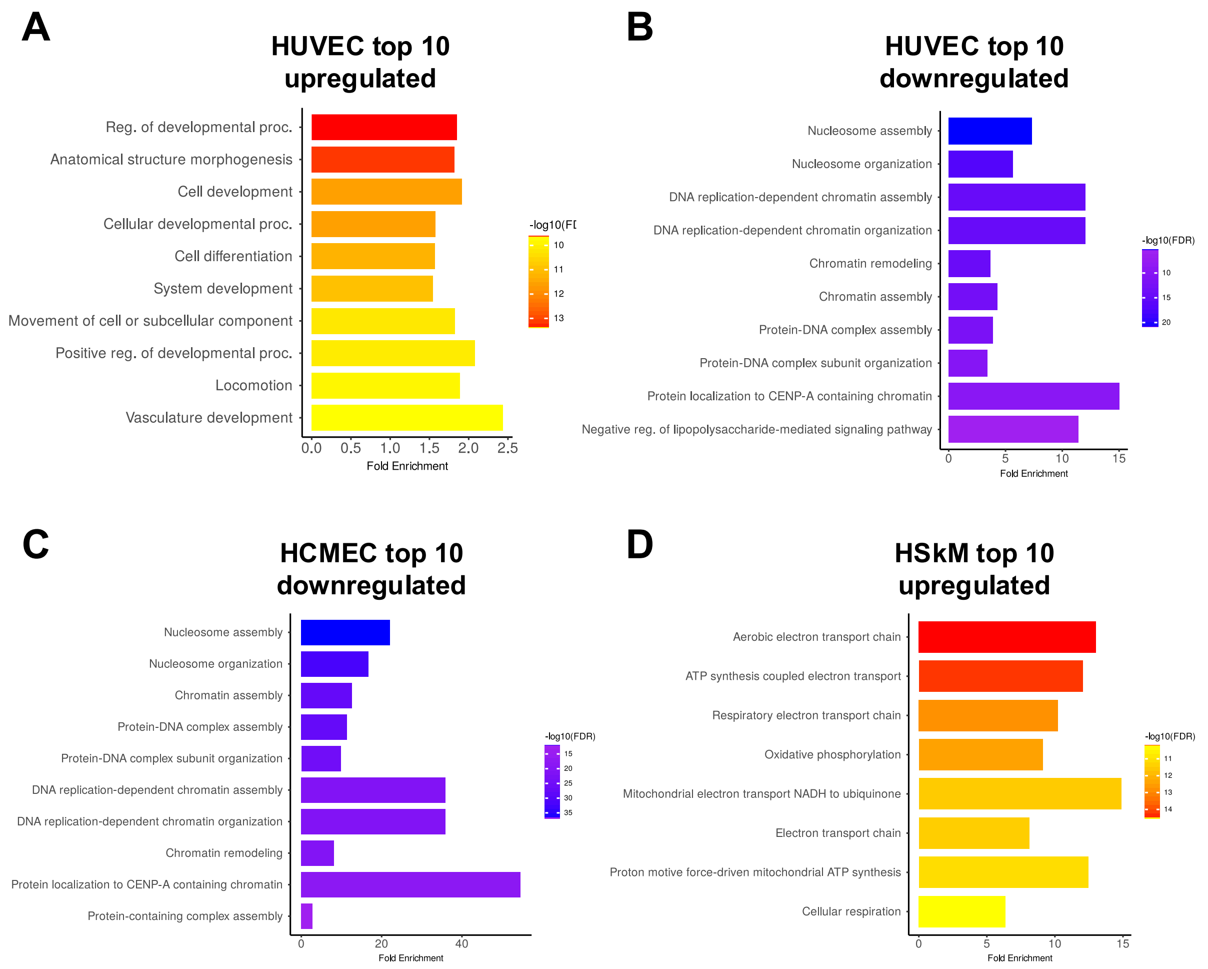


**Supplementary Fig. 2.** Gene set enrichment of GO:BP terms from overrepresentation analysis with ShinyGO web-application. Analysis was done by comparing significantly upregulated or downregulated genes to background (all expressed genes in samples after filtering) **A)** HUVEC top 10 enriched pathways from upregulated genes. **B)** HUVEC top 10 enriched pathways from downregulated genes **C)** HCMEC top 10 enriched pathways from downregulated genes. **D)** HSkM top 10 enriched pathways from upregulated genes. No significantly enriched (FDR ≤ 0.05) GO:BP pathways were found from upregulated HCMEC genes or downregulated HSkM genes. Results were visualized by the ShinyGO web-application

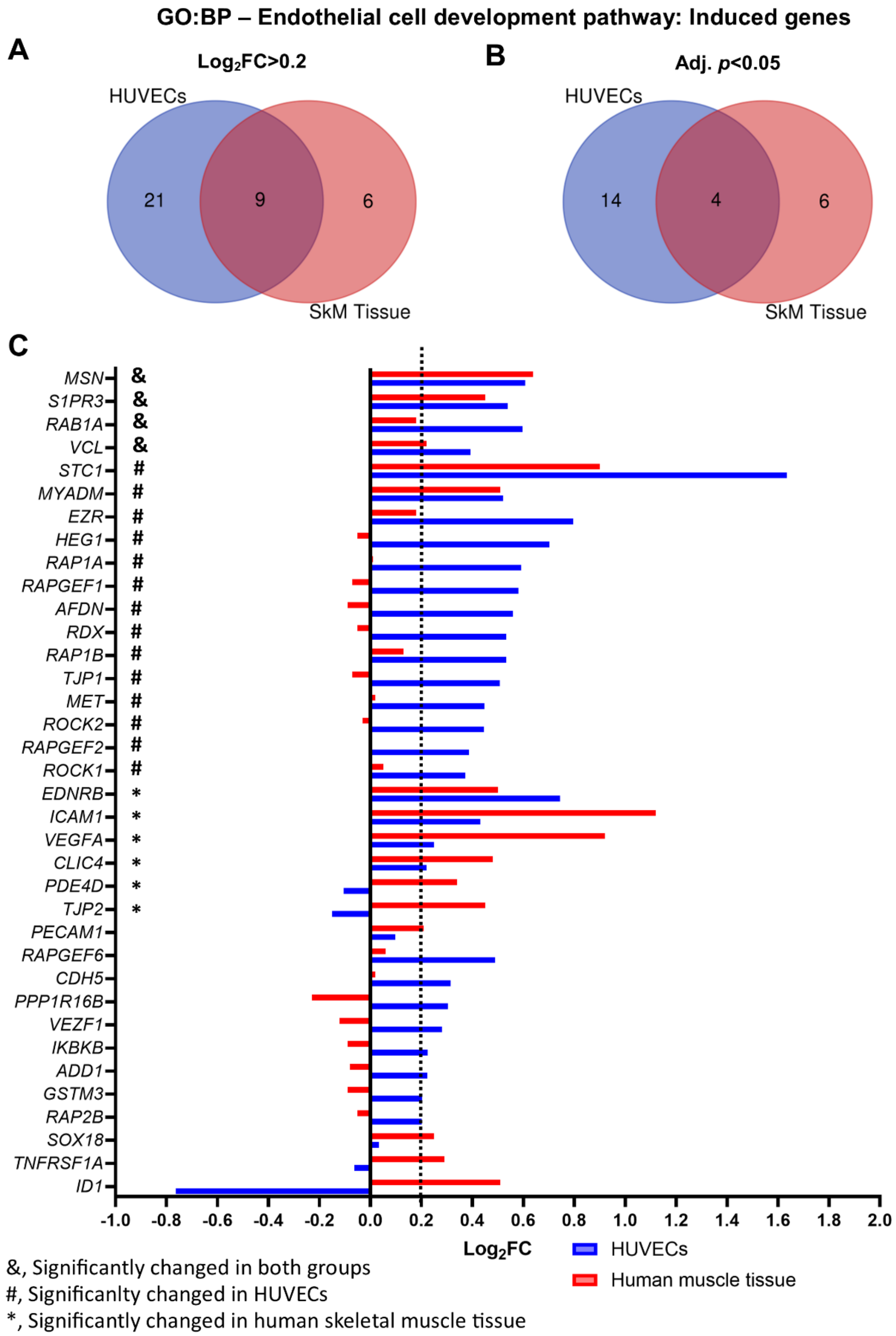


**Supplementary Fig. 3**. Induction of genes of the *Endothelial cell development* pathway (GO:0001885) in response to mechanical loading in HUVECs and acute aerobic exercise in human skeletal muscle (HSkM) tissue. **A)** A Venn diagram of the induced genes. **B)** A Venn diagram of significantly (adj. $p$<0.05) upregulated genes. **C)** Graphical presentation of mean fold changes of the genes (out of 72 genes of the pathway) with $\log_2$FC>0.2 in HUVECs or HSkM tissue. The gene expression changes in response to acute aerobic exercise were checked from MetaMEx (https://www.metamex.eu/app/metamex, health status: healthy). The Venn diagrams were prepared with a web application (https://bioinformatics.psb.ugent.be/webtools/Venn/).

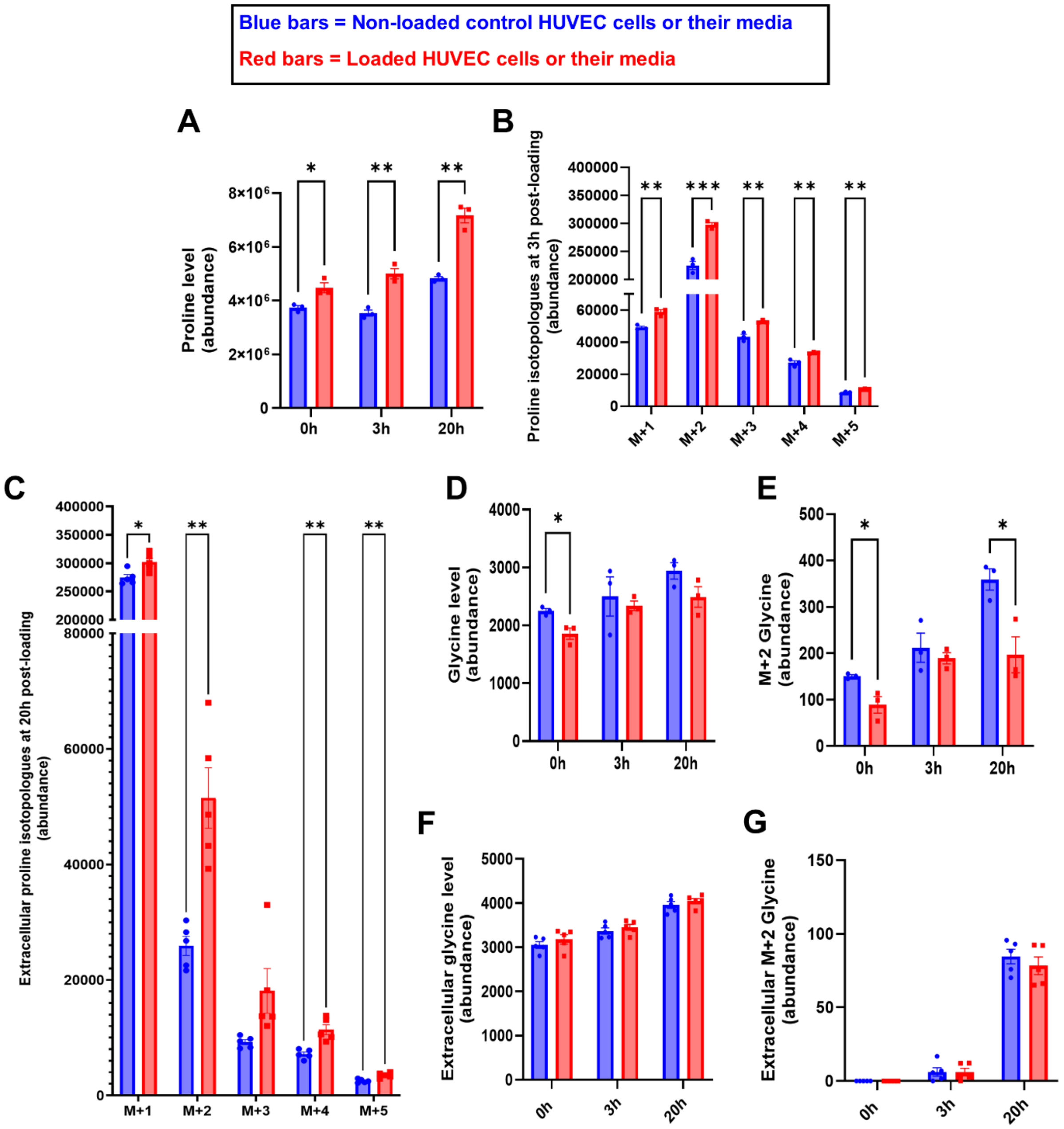


**Supplementary Fig. 4.** $^{13}C$-(U)-glucose-derived carbon incorporation into alanine, proline, and glycine in non-loaded (blue bars) and mechanically loaded (red bars) HUVECs at different time points after 5h mechanical loading.

**A)** Intracellular proline level (unlabeled + labeled).
**B)** Intracellular proline M+1-5 isotopologues at 3h post-loading.
**C)** Extracellular proline M+1-5 isotopologues at 20h post-loading.
**D)** Intracellular glycine level (unlabeled+labeled) at different time points.
**E)** Intracellular labeled (M+2) glycine level at different time points.
**F)** Extracellular glycine level (unlabeled+labeled) at different time points.
**G**) Extracellular labeled (M+2) glycine level at different time points.
Individual values are plotted, and data are presented as mean ± SEM.
**P < 0.05, **P < 0.01, ***P < 0.001* (Student's t-test).

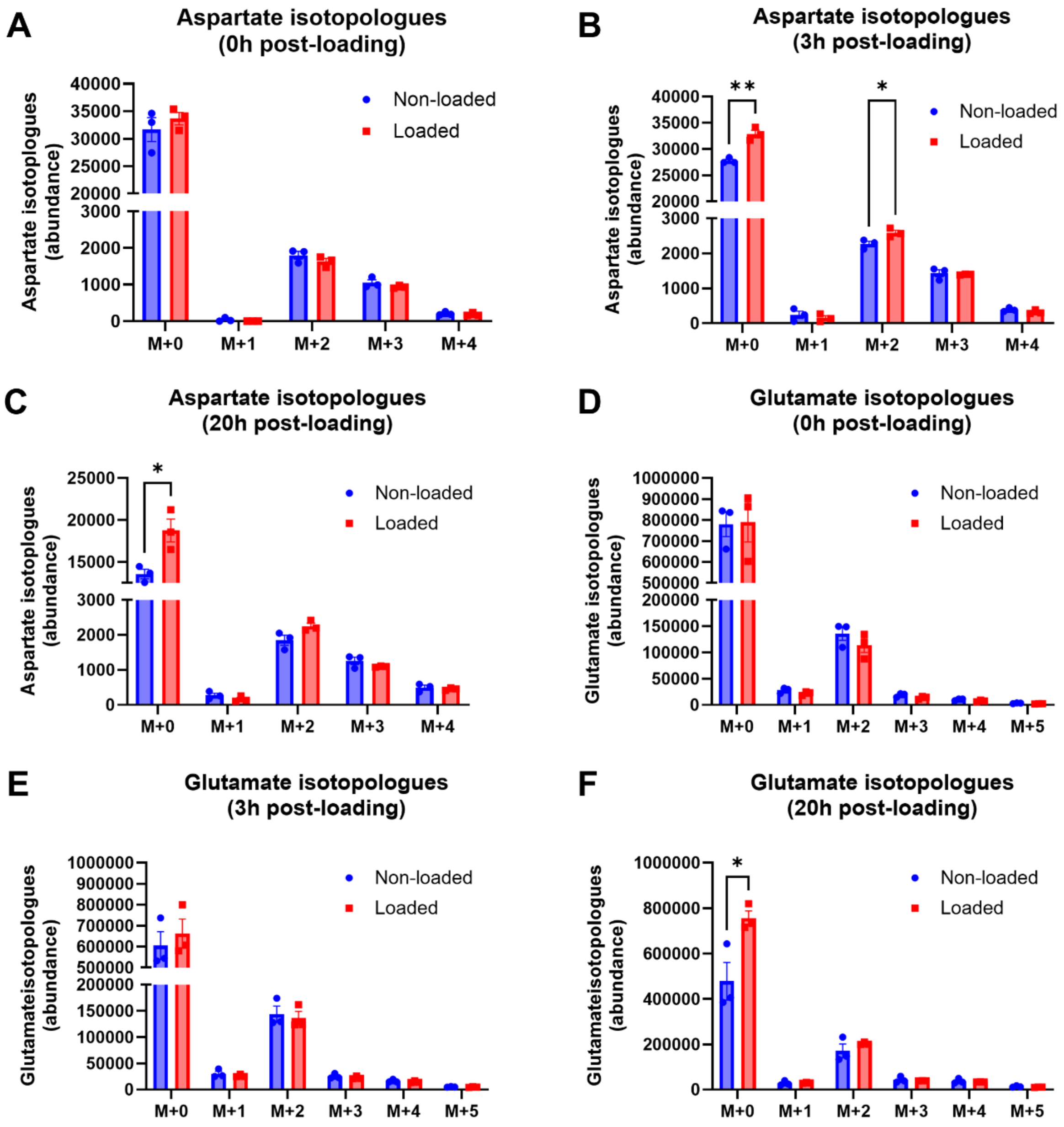


**Supplementary Fig. 5.** Labeling patterns of intracellular aspartate and glutamate in non-loaded control and mechanically loaded (5h loading) HUVECs fed with $^{13}$C-(U)-glucose.

**A)** Aspartate isotopologues at 0h post-loading.
**B)** Aspartate isotopologues at 3h post-loading.
**C)** Aspartate isotopologues at 20h post-loading
**D)** Glutamate isotopologues at 0h post-loading.
**E)** Glutamate isotopologues at 3h post-loading.
**F)** Glutamate isotopologues at 20h post-loading.
Individual values are plotted, and data are presented as mean ± SEM.
*$*P < 0.05$, $**P < 0.01$*(Student's t-test).

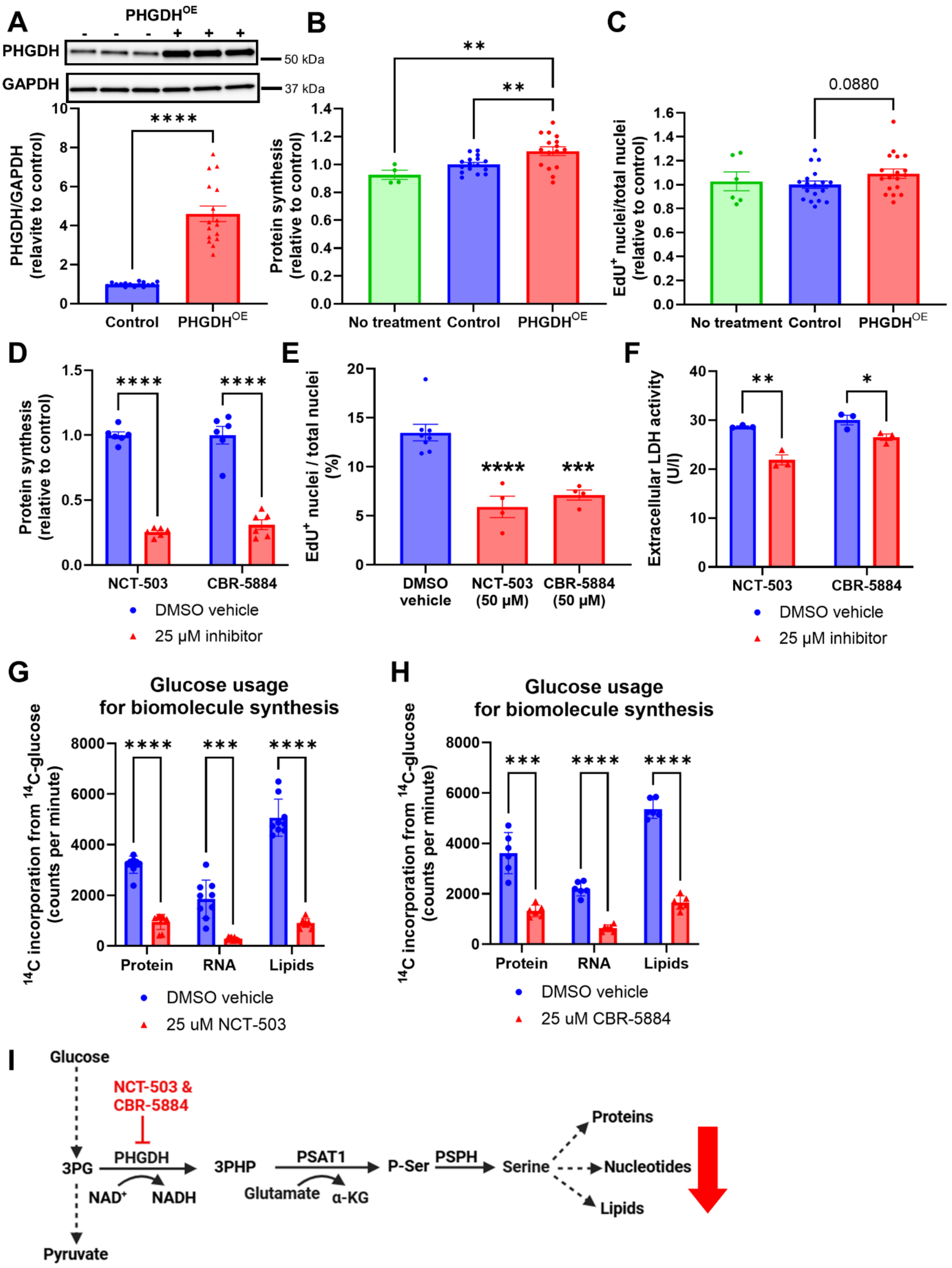


**Supplementary Fig. 6.** Effects of the serine synthesis pathway enzyme phosphoglycerate dehydrogenase (PHGDH) overexpression and PHGDH inhibitors on anabolism of HUVECs.

**A)** PHGDH level in response to overexpression treatment (n = 16).

**B)** Protein synthesis ($^{14}$C-phenylalanine incorporation into proteins) in response to PHGDH overexpression treatment (n = 16, but in no treatment group n = 4).

**C)** Cell proliferation in response to PHGDH overexpression treatment (n = 24, but in the no-treatment group, n = 6).
**D)** protein synthesis (relative $^{14}$C-valine incorporation into proteins) in response to PHGDH inhibitors NCT-503 and CBR-5884 (n = 6).
**E)** PHGDH inhibitors decreased the proliferation of HUVECs, as assessed by EdU analysis. The groups were statistically compared only to the DMSO vehicle control (n = 8 in the DMSO control group and n = 4 in the inhibitor groups).
**F)** PHGDH inhibitors did not induce cell death according to the lactate dehydrogenase (LDH) activity measured from conditioned media (i.e., increased LDH activity would indicate membrane damage and cell death) (n = 3). Individual values are plotted, and data are presented as mean ± SEM.
**G)** $^{14}$C-U-glucose-derived carbon incorporation into proteins, RNA, and lipids in response to NCT-503 (n = 9).
**H)** $^{14}$C-U-glucose-derived carbon incorporation into proteins, RNA, and lipids in response to CBR-5884 (n = 6).
**I)** Schematic presentation of the effects of PHGDH inhibitors.

Individual values are plotted, and data are presented as mean ± SEM. ****$p < 0.001$, **** $p <$ 0.0001. Student's t-test was used when two groups were compared and 1-ANOVA with Fisher's LSD test when more than two groups were compared (panels B, C & E).
